# Right posterior theta facilitates memory encoding and recall during virtual navigation

**DOI:** 10.64898/2026.02.24.707786

**Authors:** Malte R. Güth, Travis E. Baker

**Affiliations:** Center for Molecular and Behavioral Neuroscience, Rutgers University; Graduate Program in Neuroscience, Rutgers University, Newark, USA

## Abstract

Despite decades of animal and human intracranial work highlighting the critical role theta oscillations (4-12 Hz) play in memory encoding and recall stage during navigation, the link between scalp recorded human theta oscillations and spatial encoding and recall is lacking. In the present study, we used right posterior theta (RPT) – a scalp-level theta signal believed to be generated in the medial temporal cortex – to examine spatial encoding and recall during virtual spatial navigation. In particular, we recorded EEG from 27 healthy subjects performing a novel virtual Linear Track Memory (LTM) task. During the encoding stage of the task, a reward cue was presented at one of five pillar locations along a linear track. During the recall stage, subjects were presented with images of the five pillars and five new pillars, and were asked to press a button when the rewarded target pillar location appeared. If correct, subjects received 5 cents for that trial. Memory performance was assessed using reaction time, d-prime (d’), and response bias (β), and RPT was measured following the onset of the reward cue at bilateral scalp electrodes P7 and P8. Consistent with previous work, RPT peaked approximately 170-300 ms over the right hemisphere (P8) after cue onset, which was significantly increased for reward cues during the encoding stage and for the target pillar during the recall stage. Importantly, general linear model regressions revealed that peak RPT power during the encoding stage significantly predicted higher d’ and β scores during recall, supporting the relationship between RPT peak power and memory performance. Together, these findings support the proposal that RPT activity reflects the encoding of salient information for the purpose of spatial navigation and a promising candidate biomarker for memory-related functioning in health and disease (e.g., Alzheimer’s disease).

## Introduction

Theta oscillations (4-12 Hz) recorded in the hippocampus and parahippocampal gyrus (PHG) play an important role in the encoding and recall of motivationally salient information (e.g., landmarks and rewards) during navigation (Buzsáki, 2005; Buzsáki & Moser, 2013; Hasselmo & Stern, 2014). When animals encounter salient events in their environment, the precise firing rates of hippocampal place cells and parahippocampal grid cells shifts with respect to the phase of the theta rhythm and the phase is reset, which facilitates the encoding of salient information (Burgess, 2008; Givens, 1996; McCartney et al., 2004; Williams & Givens, 2003). In humans, evidence from intracranial electroencephalography (iEEG) recordings of spikes and local field potentials (LFP) has demonstrated that the amplitude and phase of hippocampus (Ekstrom et al., 2005) and PHG (Nadasdy et al., 2017; Qasim et al., 2019; Stangl et al., 2021) theta activity is sensitive to spatial features such as grid-like representations of virtual environments (Jacobs et al., 2013; Nadasdy et al., 2017), the direction of movement (Jacobs et al., 2010), velocity (Goyal et al., 2020), and spatial boundary locations relevant for goal attainment (Stangl et al., 2021). Further, iEEG studies recording from epilepsy patients have identified hippocampal and entorhinal theta oscillations tracking movement during virtual navigation (Aghajan et al., 2017; Goyal et al., 2020) and population firing rates encoding memory traces of object-location associations (Qasim et al., 2019). More specifically, Qasim et al. (2019) employed a virtual navigation task with distinct encoding and recall trials where subjects first learned the locations of objects along a linear track and then were asked to recall the locations after the objects were removed. In the entorhinal cortex, the authors found cells with firing rates that exhibited spatial tuning to specific locations containing spatial cues for rewards along the track. Importantly, these same cells during a delay period preceding recall showed similar activity patterns, which was not observed during the recall stage or in cells that lacked spatial tuning. Hence, this task engaged memory-trace cells related to the encoding of reward-related locations.

While animal and human iEEG data from epilepsy patients are highly complementary thus far, the functional role of theta oscillations during navigation in healthy human subjects remains elusive due to the limited options for using invasive recording techniques and poor spectral resolution of functional neuroimaging (fMRI, NIRS, PET). To overcome this limitation, we collected scalp-recorded EEG activity during a memory encoding and recall virtual navigation task. In line with the task used by the aforementioned iEEG studies (Goyal et al., 2020; Qasim et al., 2019), we developed an EEG-based Linear Track Memory (LTM) task consisting of an encoding and recall stage. In brief, during the encoding stage of the task, a reward cue was presented at one of five pillar locations along a virtual linear track. During the recall stage, subjects were presented with images of the five pillars, plus five new pillars, and were asked to identify the location of the reward cue by pressing a button. If correct, subjects received a reward for the trial. In regards to the electrophysiological data, we focused our analysis on right posterior theta (RPT), an electrophysiological signal sensitive to salient events encountered during virtual navigation (Baker & Holroyd, 2009, 2013). RPT describes a pattern of phase reset and power enhancement in right posterior electrodes (5-10 Hz, 170–300 ms) found to be sensitive to the spatial position of reward feedback during virtual navigation. We have proposed that RPT reflects partial phase resetting (i.e., concomitant increases in phase coherence and spectral power relative to baseline) of the ongoing theta rhythm in the PHG, a mechanism thought to facilitate the encoding of motivationally relevant spatial information for navigation (Güth et al., 2025). This phenomenon is also observed in the time domain as an event-related potential (ERP) called the topographical N170 (Baker & Holroyd, 2009) and is sensitive to individual differences in spatial ability (e.g., how well subjects recalled the shape of the maze they had navigated; Baker et al., 2015; Baker & Holroyd, 2013)

Converging evidence across multiple neuroimaging modalities supports the PHG as the source of RPT. Asynchronous EEG-MEG recordings demonstrated that the spatial sensitivity of RPT (earlier and larger responses for right vs. left alleys) was reproducible across modalities, with MEG data ruling out volume conduction artifacts (Güth et al., 2025). Consistent with these findings, independent fMRI data showed greater right PHG activation during the same virtual navigation task compared to a non-spatial control condition, with preferential activation for right relative to left alleys (Baker et al., 2015). Most compellingly, simultaneous EEG-fMRI recordings revealed that single-trial evoked (i.e., phase-locked) RPT power (7-8 Hz) significantly predicted single-trial hemodynamic responses in both posterior and anterior PHG clusters (Güth et al., 2025), directly linking scalp-recorded theta oscillations to PHG hemodynamic activity during goal-directed navigation.

While these findings indicate that feedback processing during virtual navigation can elicit phased-locked RPT responses, whether or not these signals reflect the encoding of the event for the purpose of memory formation remains unknown. Thus, our purpose was twofold: First, given the novelty of the task, we tested whether the cues presented in the LTM task could elicit RPT. Second, if RPT is in fact associated with memory encoding and recall, individual variation in RPT power during the encoding stage should predict individual variation in memory performance during the recall stage such as d-prime (d’). In summary, the present study provides support for the proposal that RPT is associated with parahippocampal processes related to the encoding of salient information for the purpose of spatial memory and navigation.

## Methods

### Participants

31 undergraduate students (M_Age_ = 25 ± 2.76 years; 5 female) were recruited from Rutgers University Newark and the New Jersey Institute of Technology. They were compensated with course credit as well as a monetary bonus equivalent to their memory performance in the LTM. Four subjects were excluded due to poor EEG data quality and a low number of usable trials, bringing the final sample to 27 subjects (M_Age_ = 24.8 ± 3 years; 5 female). All participants were right-handed (18-35 years old), gave informed consent, had normal or corrected-to-normal vision, no history of neurological or psychiatric disorders or head trauma, and completed self-report questionnaires assessing demographics, handedness (Oldfield, 1971), and spatial navigation ability (Hegarty et al., 2002). The study was approved by the Rutgers University Ethics Committee Board and was conducted in accordance with the ethical standards prescribed in the 1964 Declaration of Helsinki.

### Linear Track Memory Task

The virtual LTM task was constructed using commercially available 3D modeling software (Home design 3D; Expert Software Inc., Coral Gables, FL). Each trial of the LTM task was divided into two stages: an encoding stage (Figure 1A and 1B) and a recall stage (Figure 1C). During the encoding stage subjects were placed at the start of the track and following a go signal, subjects traversed the linear track towards a brick wall while passing five pairs of pillars with distinct textures and colors. Pillars were placed to both the left and the right side of the track. When subjects reached each pillar pair location (ISI between pillars: 400 ms), they stopped for 150 ms before an apple (reward cue) or an orange (no reward cue) appeared in the center of the track for 350 ms. On each trial only one apple was shown at a random pillar location while an orange appeared at the remaining pillar locations. Alternatively, on 16.6% of the trials none of the pillars had an apple. After reaching the wall at the end of the track, the recall stage started. In this stage subjects were shown a list of 10 pillars in one of three pseudo-randomized order sequences (250 ms ISI between response and next pillar), including the five pillars from the track and five distractor pillars. Subjects were instructed to indicate via button press which of these pillars was associated with an apple without a time limit and were told that they would increase their monetary bonus by correctly identifying the reward pillar (5 cents per correct response). For responding, subjects were instructed to press the left button on the response box with their left index finger to identify a pillar as a reward pillar (apple) and the right button with their right index finger for all no reward pillars (orange) and distractors. In total, subjects performed 120 trials across four blocks (30 trials each), with each trial lasting approximately 12 seconds. Participants viewed all stimuli from a distance of approximately 70 cm (13.9° wide, 9.8° high) and displayed on a 24-inch computer monitor using E-Prime experiment control software (Psychological Software Tools, Pittsburgh, PA).

**Figure 1.**
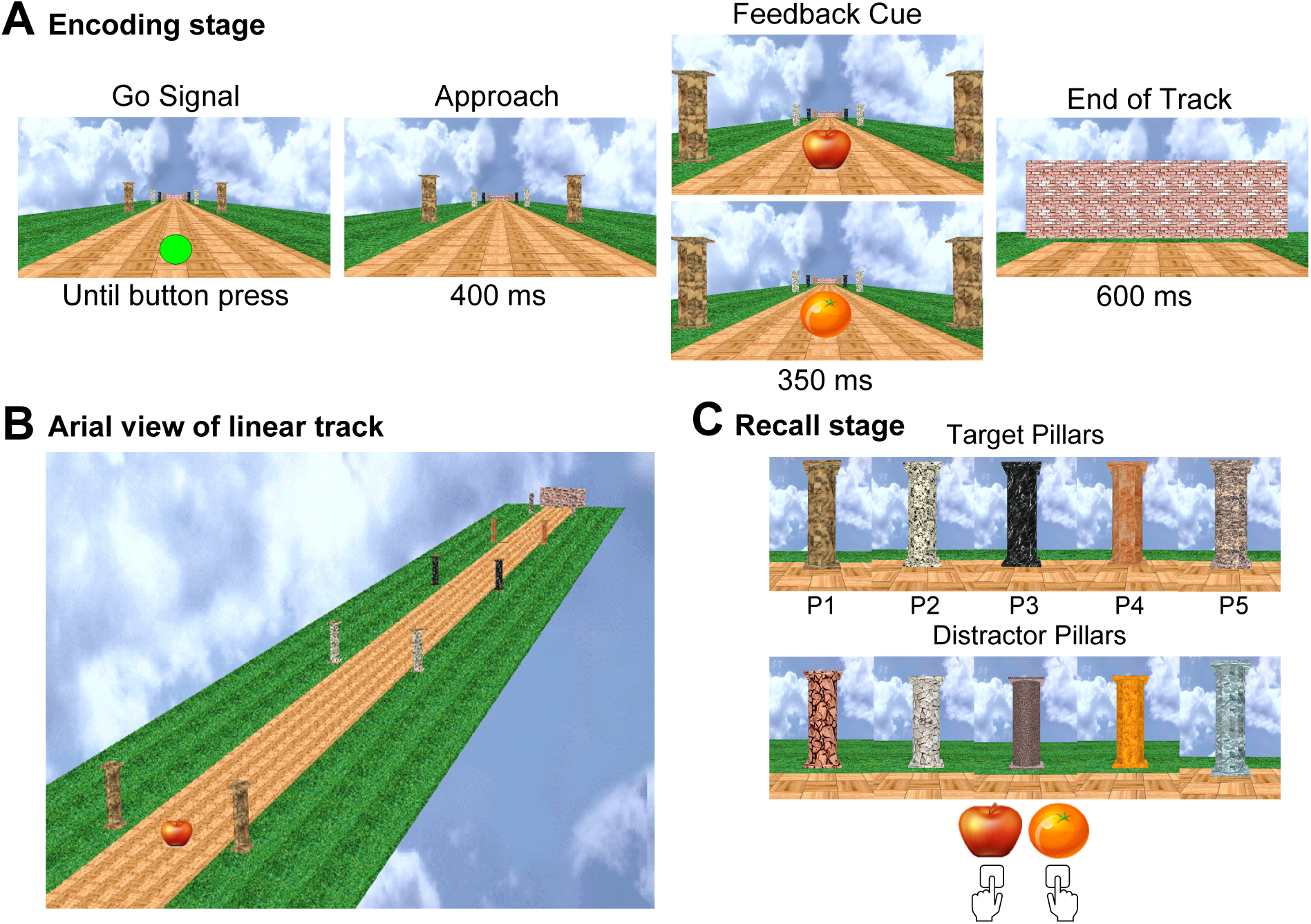
Linear Track Memory task. **A**: Main events on a LTM trial. Subjects started at the beginning of the track and received a go signal (button press to start, self-paced), before moving to each pillar pair and encountering the feedback cue (apple or orange). Each pillar pair was made distinct through its texture and color. When subjects finished the track (End of Track), they entered the recall stage. **B**: Overview of the encoding segment of the LTM task, starting with feedback at the first pillar pair (bottom left). **C:** Pillar displays shown to subjects during the recall stage. The top row shows the potential target pillars (P1-P5) and the bottom row the distractor pillars. Subjects indicated via button press which pillar marked a reward cue position on the track (left index finger for apple pillar, right index finger for all others). Pillars were presented in one of three pseudo-randomized order sequences.

### Behavioral analysis

All behavioral analyses were focused on the recall stage. Responses during the recall stage were categorized as hits (correct identifications of target pillars), misses (missed target pillars), correct rejections (correct rejections of non-target pillars and distractors), and false alarms (incorrect identifications of non-target pillars and distractors as target pillars). Memory performance was assessed using hit rates (ratio of hits divided by sum of hits and misses), false alarm rates (ratio of false alarms divided by the sum of false alarms and correct rejections), and d’, which calculates discriminability as the standardized difference between the hit rate and false alarm rate distributions (Green & Swets, 1974; Macmillan & Creelman, 1990). Higher d’ indicate that subjects were better able to distinguish between target and non-target pillars. To quantify participants’ response bias, the parameter β was calculated using the difference between the z-scored, squared false alarm and hit rates. This difference was then converted using an exponential transformation. β reflects the participant’s tendency to adopt a more liberal (lower β value) or conservative (higher β value) decision criterion when classifying memory items. As a result, higher values indicate more cautious responding minimizing false positives, while lower values indicate higher readiness to identify targets minimizing misses.

To further investigate how confident subjects were in their response, reaction times (RT) for all target and distractor pillars during the recall stage were analyzed. For the analysis of RT, responses below 100 ms and above 2500 ms were excluded. Performance measures were compared across the levels of the factor Pillar Location (positions 1-5) in one-way repeated measures ANOVAs for d’ and RT. Paired t-tests were run for post-hoc comparisons and p-values were corrected for multiple comparisons using the Bonferroni-Holm method. To test the homogeneity of variance and approximate normal distribution of the residuals, Levene tests and a Shapiro-Wilk tests were run. If the former indicated a violation of the sphericity assumption, the degrees of freedom were adjusted using the Greenhouse-Geisser method (Greenhouse & Geisser, 1959). All statistical analyses were run in Python 3.10 using the scipy (Virtanen et al., 2020), pingouin (Vallat, 2018), and statsmodels packages (Seabold & Perktold, 2010) as well as the rstatix package in R (R Development Core Team, 2016).

### EEG data acquisition

EEG data were recorded at 32 electrodes in accordance with the extended international 10-20 system (Jasper, 1958) using Ag/AgCl ring electrodes mounted in a nylon electrode cap with an abrasive, conductive gel (Falk Minow Services, Herrsching). In addition, two electrodes were placed on the left and right mastoids, and one was placed above the right eye. These channels were amplified by low-noise electrode differential amplifiers with a frequency response of DC 0.017-67.5 Hz (90 dB octave roll off). Signals were then digitized at a sampling rate of 1000 Hz, online referenced to an average of all channels, and were recorded using Brain Vision Recorder software (Brain Products GmbH, Gilching, Germany). Before the experiment all inter-electrode impedances were reduced to a value below 20 kΩ.

### EEG data preprocessing

EEG analyses were performed in Python 3.10 and MNE-python (Version 1.0.0) for M/EEG analysis (Gramfort et al., 2014). Data were bandpass filtered (low-pass: 60 Hz, high-pass: 0.1 Hz) to remove power line noise, other high frequency noise and slow, movement induced signals. To remove ocular and movement artifacts, data were entered into an Independent Component Analysis (ICA) and decomposed into independent components using the Infomax algorithm. The ICA was run on a copy of the data, which was high-pass filtered at 1 Hz to prevent distortion through high amplitude, low frequency signals (Winkler et al., 2015). For the same reason, excessively noisy channels were interpolated before the ICA. Next, filtered EOG signals were segmented around likely saccades and blinks. The time courses of each component were correlated to these ocular artifact segments in order to identify components representing artifact signals. Furthermore, potential artifact components were inspected based on their topography and spectral composition. Components marked as artifacts were excluded from back-projection to continuous data. The cleaned EEG data were re-referenced to the average of all sensors. Lastly, all data were segmented around feedback cue presentation at each pillar (±2500 ms) for the encoding stage and around target and non-target pillars for the recall stage. Segments exceeding a maximum peak-to-peak amplitude of 150 μV were marked and excluded from the time-frequency analysis.

### Time-frequency analysis

For the time-frequency analysis, cleaned segments time-locked to feedback presentation for each pillar, were exported to MATLAB (release 2020b, Mathworks, Massachusetts, USA) and analyzed using custom MATLAB scripts from our previous studies (Baker & Holroyd, 2013). The frequencies ranging from 1-50 Hz were analyzed using a complex seven-cycle Morlet wavelet for convolution. Total spectral power was obtained by averaging the EEG spectrum across all trials and time for each subject. Time-frequency analysis on the single trial EEG data thus yielded total theta power, including theta power that was both phase consistent (evoked) and phase inconsistent (induced) across trials with respect to the eliciting event. The relative change in the power for each pillar location was determined by averaging the baseline activity (−200 to −100 ms prestimulus) across time for each frequency and then subtracting the average from each data point following stimulus presentation for the corresponding frequency. This value was then divided by the baseline activity to normalize the change of power to the baseline activity. To note, spectral power was reported unitless because it is calculated as a proportional increase/decrease relative to baseline. The peak RPT power and latency were obtained by averaging power values between 5-8 Hz and detecting the maximum power between 100 ms and 600 ms following the onset of the cue stimulus, which is consistent with the frequency range associated with trial-by-trial PHG activation (Güth et al., 2025).

Total RPT values recorded during the encoding stage were then analyzed using a three-way repeated measures ANOVAs including the factors Channel (P7, P8), Cue Valence (reward cue, no reward cue), and Pillar Location (P1, P2, P3, P4, P5). For the recall stage, another three-way repeated measures ANOVA was run with the factor Pillar Type instead of Cue Valence. Pillar Type included the factor levels target pillar, which denoted the pillar associated with reward cue in the previous encoding stage, and non-target pillar (pillars paired with no reward cues). In addition, a two-way repeated measures ANOVA focused on comparing hits and misses using the factor Response Type as well as the encoding and recall stage using the factor Task Stage. Post-hoc paired t-tests were run to compare factor levels of significant effects and the p-values were corrected for multiple comparisons using the Bonferroni-Holm method. Finally, general linear models (GLM) predicting memory measures (d’, β, RT) were fit with parametric regressors based on z-standardized RPT power recorded from P8 during the encoding and recall stage to test whether RPT is connected to successful memory encoding or recall. All statistical analyses of time-frequency power were run using the same software environments and packages as for the behavioral analysis.

### Data and code availability

Analysis scripts, documentations, and examples for both pre-processing and higher level statistics can be found in the github repository for this study: https://github.com/BakerlabRutgers/ltm. All further code and data are available upon reasonable request.

## Results

### Memory performance

On average, subjects responded during the recall stage with an average hit rate of 87% (SD = 15%) and an average d’ score of 2.49 (SD = 0.65) across all trials. A one-way repeated measures ANOVA on d’ score with Pillar Location (P1, P2, P3, P4, P5) as a within-group factor yielded a significant main effect, 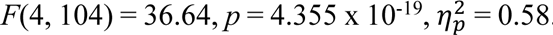. Post-hoc tests revealed that P3 (M = 3.81, SD = 0.86) had significantly higher d’ values than P1 (M = 1.83, SD = 0.7; *t*(26) = 15.23, *p* = 1.812 x 10^-13^, *d* = 2.98), P2 (M = 3.25, SD = 0.97; *t*(26) = 3.39, *p* = .008, *d* = 0.66), P4 (M = 2.98, SD = 0.77; *t*(26) = 5.66, *p =* 4.142 x 10^-5^, *d* = 1.11), and P5 (M = 2.75, SD = 0.99; *t*(26) = 5.17, *p* = 1.074 x 10^-4^, *d* = 1.01, Bonferroni-Holm corrected). To note, P1 had significantly lower d’ than all other locations (*p* < .05). Figure 2A depicts differences in d’ between the five pillar locations, which followed a cubic trend. This was confirmed by fitting a mixed effects regression including both a linear (*t*(26) = 0.74, *b* = −0.64, *p* = .461) and a cubic term (*t*(26) = −2.1, *b* = −0.28, *p* = .035) for Pillar Location. There was a significant, negative cubic term, indicating that performance levels started low at P1, rose towards P3 and declined again for P4 and P5. The same ANOVA was calculated with the β parameter as the dependent variable, revealing that P1 had higher β values the other pillars, indicating participants were more hesitant to identify P1 as the target (supplementary Figure S1).

**Figure 2.**
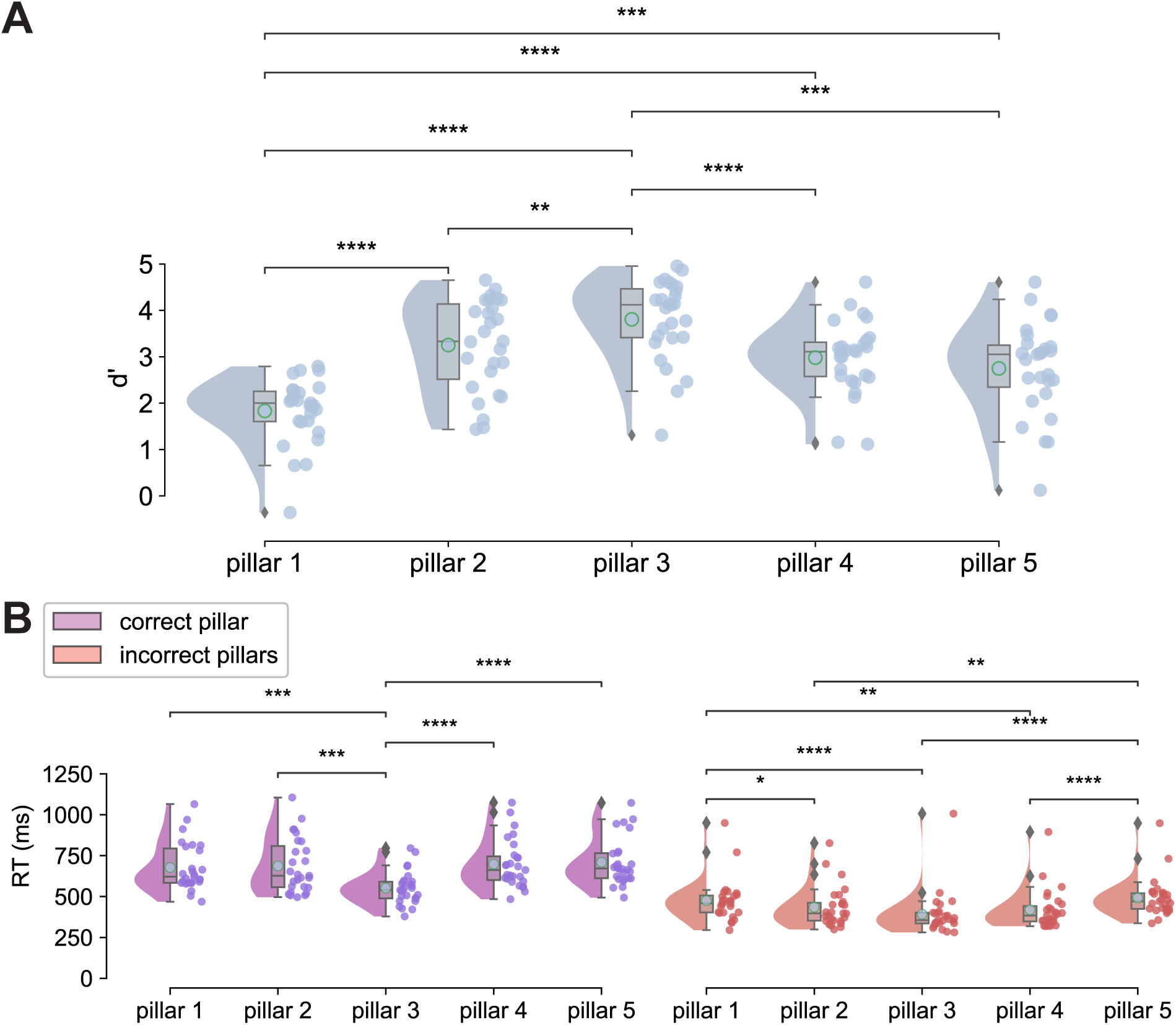
Memory performance measures. **A**: Memory performance during the recall stage as assessed by d’ for each pillar location. Raincloud plots are composed of a density plot showing the distributions d’ scores, as well as a boxplot and scatterplot showing the same data. Cyan dots in each box plot mark the mean value of each distribution and gray diamonds mark outlier values. **B**: Analogous plots to A showing RT values only for responses to correct pillars during the recall stage (purple) and RT values only for incorrect pillars (orange). *\*p < .05, **p < .01*, \*\*\**p < .001*, \*\*\*\**p < .0001*

Next, a one-way repeated measures ANOVA on correct recall RT with Pillar Location (P1, P2, P3, P4, P5) as a within-subjects factor (Figure 2B) revealed a significant main effect, 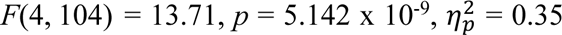 When P3 contained the reward cue (M = 552 ms, SD = 98 ms), it was identified as the correct location significantly faster than P1 (M = 678 ms, SD = 144 ms; *t*(26) = 4.91, *p =* 3.388 x 10^-4^, *d* = 0.96), P2 (M = 687 ms, SD = 165 ms; *t*(26) = 4.48, *p =* 9.312 x 10^-4^, *d* = 0.88), P4 (M = 698 ms, SD = 145 ms; *t*(26) = 7.34, *p* = 8.579 x 10^-7^, *d* = 1.44), and P5 (M = 710 ms, SD = 142 ms; *t*(26) = 6.23, *p* = 1.23 x 10^-5^, *d* = 1.22). Finally, when testing RT for incorrect pillar locations, a significant main effect of Pillar Location was observed, 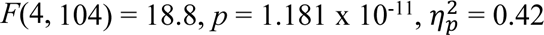. There were lower RT values for P3 (M = 404 ms, SD = 137 ms) compared to P1 (M = 490 ms, SD = 132 ms; *t*(26) = 5.91, *p* = 2.508 x 10^-5^, *d* = 1.16) and P5 (M = 507 ms, SD = 122 ms; *t*(26) = 7.91, *p* = 2.209 x 10^-7^, *d* = 1.55). P5 also had slower RT than P4 and P2 (*p* < .05, Bonferroni-Holm corrected).

### EEG results

#### Encoding Stage

Figure 3 depicts EEG results time-locked to the start of the track for both reward (apples) and no reward cues (oranges). In the time-frequency domain (Figure 3A), a large increase in total RPT power was observed between 4-12 Hz following the onset of the cues at one of the five pillar locations, which was most pronounced between 6-8 Hz at P8 and peaked at approximately 258 ms (SD = 81 ms) after each cue presentation. A three-way repeated measures ANOVA on total RPT power with Pillar Location (P1, P2, P3, P4, P5), Cue Valence (reward cue, no reward cue), and Channel (P8, P7) as within-subjects factors revealed a significant main effect of Pillar Location 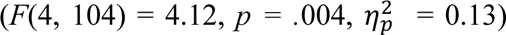. P1 (M = 1.05, SD = 0.6) elicited significantly enhanced RPT relative to P5 (M = 0.73, SD = 0.67; *t*(26) = 4.29, *p =* .005, *d* = 0.59), but not compared to P2 (M = 0.9, SD = 0.39), P3 (M = 0.91, SD = 0.56), and P4 (M = 0.97, SD = 0.52; *p* > .05). Further, a main effect of Cue Valence was also observed, 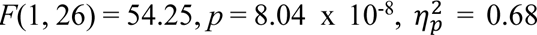, indicating greater RPT power for the reward cues (M = 1.21, SD = 0.62) compared to no reward cues (M = 0.61, SD = 0.27; Figure 3B). Lastly, a significant main effect of Channel was observed, 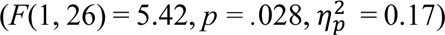, indicating stronger RPT power over channel P8 (M = 1.04, SD = 0.55) compared to channel P7 (M = 0.78, SD = 0.48). Finally, there was a significant interaction of the factors Valence and Channel 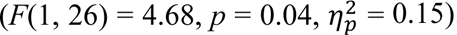. RPT power for reward cues was significantly higher at P8 (M = 1.38, SD = 0.73) than at P7 (M = 1.04, SD = 0.66; *t*(26) = 2.5, *p =* .038, *d* = 0.49), but RPT power elicited by no reward cues was the same at P8 (M = 0.7, SD = 0.39) and P7 (M = 0.52, SD = 0.34; *t*(26) = 1.9, *p =* .069, *d* = 0.37). For descriptive purposes, we present the data in the time-domain in the supplementary materials. Visual inspection of Figure S2 shows there was a robust ERP (i.e., N170) over bilateral parieto-occipital channels following the onset of feedback-related cues at each pillar location, consistent with our previous studies (see supplementary Figure S2, Baker & Holroyd, 2013; Güth et al., 2025).

**Figure 3.**
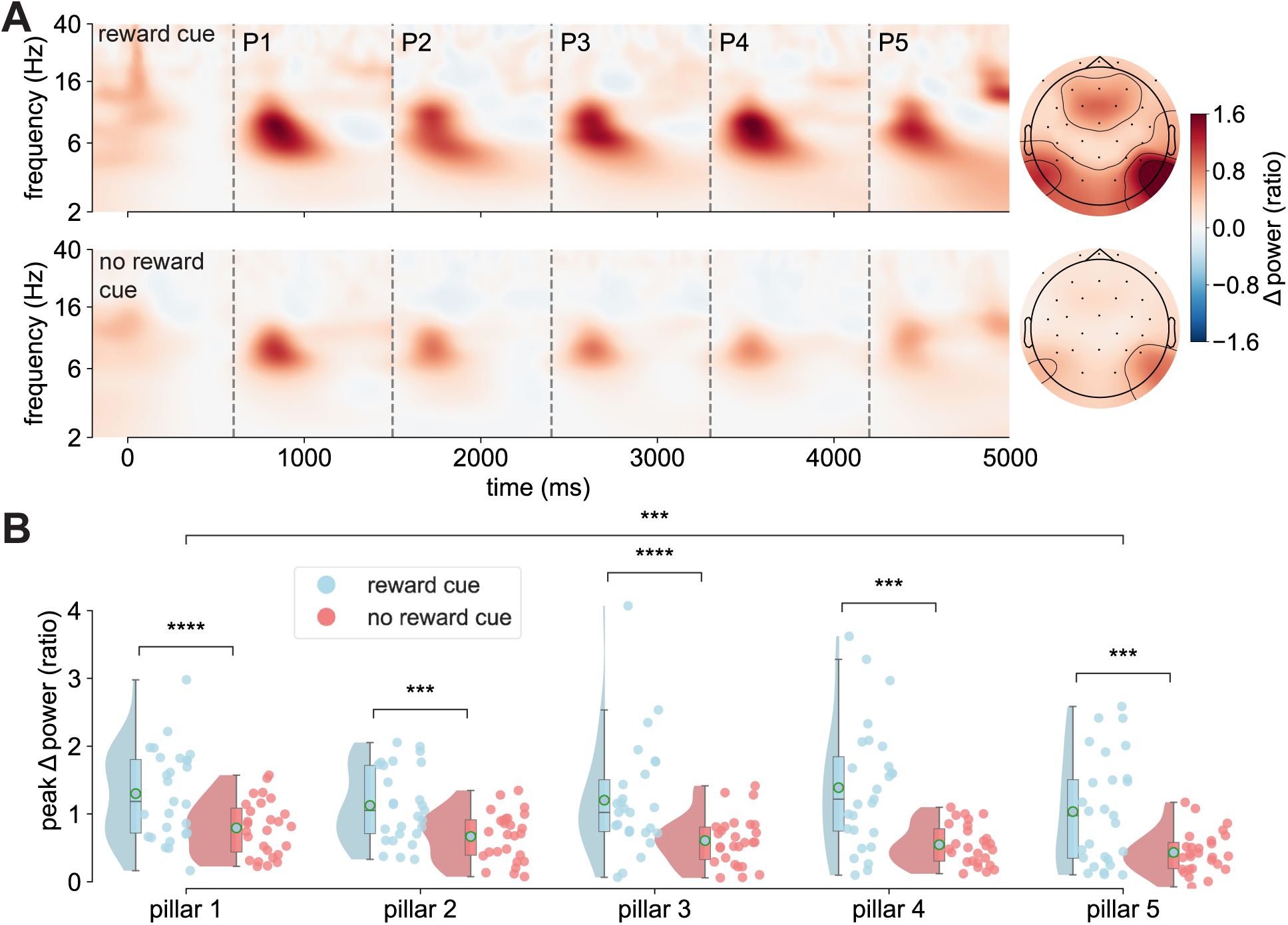
Encoding-related total RPT by cue valence. **A**: Averaged time-frequency spectrogram of the total RPT power at channel P8 for reward cues (top row) and no reward cues (bottom row). Dashed lines (P1-P5) mark the timings of cue presentations at the five pillar pairs. Topographies show the RPT power values (5-8 Hz) from 200 ms to 300 ms after reward and no reward cues averaged across the pillar locations. **B**: Peak total RPT power values (5-8 Hz) during recall stage by pillar location and valence (blue: reward cue, red: no reward cue). Values were averaged across P7 and P8. \*\*\**p < .001*, *\*\*\*\*p < .0001*

#### Recall Stage

Analyzing the time-frequency data time-locked to target and non-target pillar presentations in the recall stage also revealed strong increases in total RPT power (Figure 4A). The same three-way ANOVA run for the encoding stage was used for the recall stage, including the factors Pillar Location (P1, P2, P3, P4, P5), Channel (P8, P7) and Pillar Type (target, non-target pillar) instead of Cue Valence. There were both a significant main effect for Pillar Location 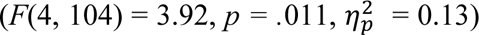 and Pillar Type 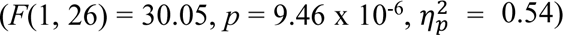. In regards to Pillar Location, P1 (M = 0.43, SD = 0.3; Figure 4B) and P2 (M = 0.44, SD = 0.38) had significantly larger RPT power than P5 (M = 0.24, SD = 0.19; P1: *t*(26) = 3.69, *p = .*01, *d* = 0.72; P2: *t*(26) = 3.2, *p = .*033, *d* = 0.63). In regards to Pillar Type, RPT was significantly enhanced in response to target pillars (M = 0.45, SD = 0.26) compared to non-target pillars (M = 0.26, SD = 0.17). There were no significant main effects or interaction effects involving the factor Channel, indicating that effects were uniform across P7 and P8. To note, visual inspection of Figure 4A reveals a large increase in beta power (13-30 Hz) within 200 ms after pillar presentation during the recall. An analysis of this beta power and cross-frequency couplings with RPT can be found in the supplementary materials (supplementary Figures S3 and S4).

**Figure 4.**
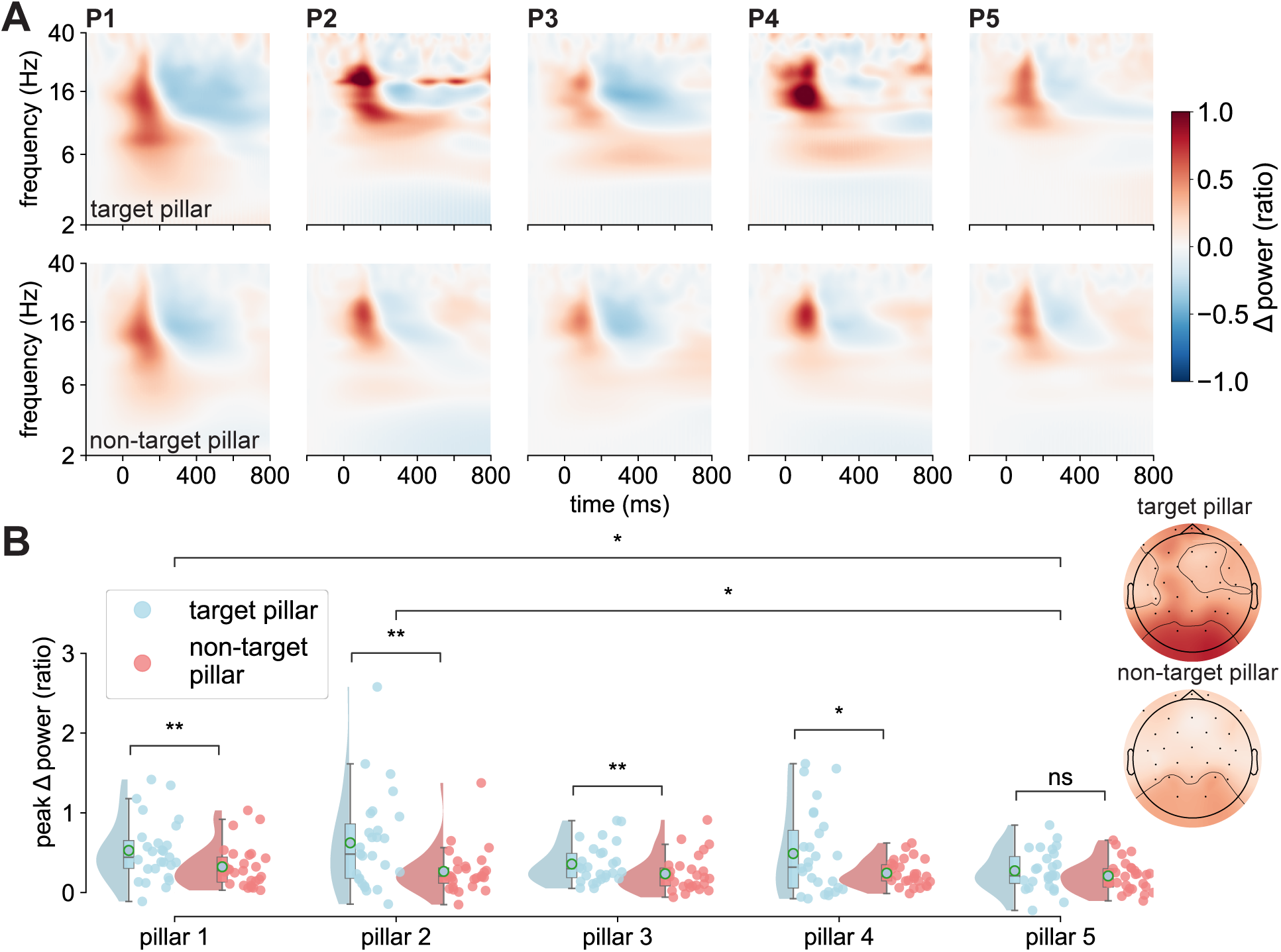
Recall related RPT by pillar type. **A**: Averaged time-frequency spectrogram of the EEG total RPT power at channel P8 time-locked to the presentation of target (top row, pillar with reward cue during encoding) and non-target pillars (bottom row, pillars with no reward cues during encoding). **B**: Peak total RPT power values (5-8 Hz) during recall stage by pillar location and pillar type averaged across P7 and P8. Topographies show the mean RPT power from 100 ms to 200 ms after pillar presentation separately for target and non-target pillars. The same color scale for RPT power values as in 4A applies. *ns: not significant, *p < .05*, *\*\*p < .01*

### Relationship of RPT and memory performance

To investigate the relationship of RPT power during the encoding/recall stage and memory performance, we tested whether there are linear relationships between RPT power and memory performance. d’ (discriminability) was used as a measure of how successful subjects differentiated between the target pillar and other pillars. GLMs predicting single subject d’ values using regressors based on the RPT power following the target pillar. Separate models for RPT during the encoding and the recall stage were fitted. During the encoding stage, single subject total RPT values significantly predicted memory performance as assessed by d’ (Figure 5A, left), *t*(26) = 2.26, *p = .*024, *b* = 0.412, *CI_95%_* = 0.055-0.769. By contrast, there was no significant relationship between RPT and d’ during the recall stage (Figure 5A, right; *t*(26) = 0.38, *p = .*702, *b* = 0.076, *CI_95%_* = −0.315-0.467). Similarly, β (response bias) was significantly predicted by RPT only during the encoding stage (Figure 5B, left; *t*(26) = 2.37, *p =* .018, *b* = 0.428, *CI_95%_* = 0.074-0.782). No such effect was found for RPT power from the recall stage (Figure 5B, right; *t*(26) = 0.17, *p =* .863, *b* = −0.035, *CI_95%_* = −0.426-0.357). Finally, we tested if high RPT power characterized the participants who scored highly in both d’ and β. For this purpose, a final GLM predicting RPT power from the encoding using regressors for d’ and β as well as their interaction term was fitted. This interaction term reached significance with a positive beta coefficient (Figure 5C, left; *t*(26) = 3.14, *p =* .002, *b* = 0.558, *CI_95%_* = 0.209-0.907), indicating that the effect of β in predicting RPT power increases as d’ increases. Therefore, RPT power during encoding was highest in participants who achieved both high d’ and β. Again, this was not the case for RPT power during recall (*t*(26) = 0.07, *p =* .942, *b* = −0.017, *CI_95%_* = −0.489-0.454; Figure 5C, right).

**Figure 5.**
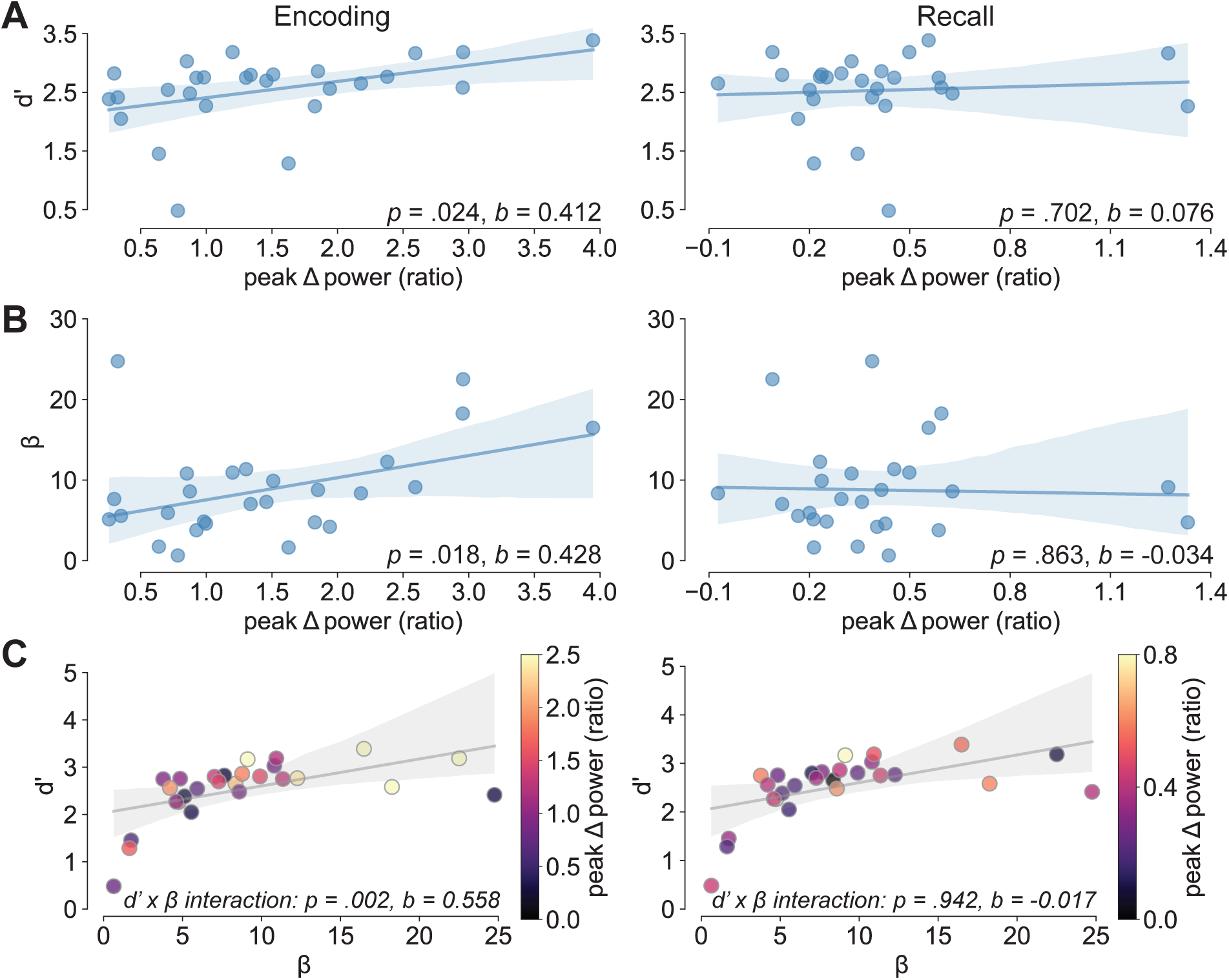
Association of memory measures and RPT. **A**: Regression lines showing the relationship between single subject d’ values and peak total RPT power for the encoding (left) and recall stage (right). Shaded areas mark a 95% confidence interval around the central tendency. **B**: Analogous plots to **A** depicting the relationship between single subject RPT and β values. **C**: Regression plots highlighting the interaction between d’ and β in predicting RPT power for the encoding stage (left) and the recall stage (right). Dots mark the d’ and β of individual subjects and are colored by the respective subject’s RPT power.

## Discussion

Animal and human studies indicate that phase resetting of theta oscillations in the PHG by salient events (e.g., rewards, landmarks) facilitates the encoding of goal-oriented information during navigation (Burgess, 2008; Givens, 1996; Hasselmo, 2008; McCartney et al., 2004; Rizzuto et al., 2003; Williams & Givens, 2003). Consistent with this work, we demonstrated that feedback cues presented in a virtual T-maze environment elicit a partial phase reset of the oscillation over right posterior electrodes. Further, the latency, power, and phase angle of RPT were found to be sensitive to feedback cues received following a right turn relative to a left turn in the T-maze (Baker & Holroyd, 2013) and single-trial RPT power during spatial navigation was linked to single-trial PHG activation (Güth et al., 2025). In line with these observations, we proposed that RPT reflects a phase-coded signal by the PHG for encoding contextual information about reward location in the environment. However, whether RPT is functionally related to memory encoding and subsequent retrieval performance remains unknown. The current study directly tests this relationship by examining whether RPT power during encoding predicts memory performance for reward-location associations.

In particular, we investigated whether scalp-recorded RPT signals during a novel virtual LTM task exhibited similar memory-related properties as found in previous human iEEG studies (Qasim et al., 2019). Following the design of Qasim et al. (2019), participants navigated a virtual linear track and learned object-reward locations during an encoding phase, then later recalled those locations during a retrieval phase. Just like in the study by Qasim et al., participants encoded one out of five possible object locations (four in Qasim et al.) and had to retrieve this location in a separate recall trial, maintaining the core encoding-retrieval structure. Overall, participants successfully learned the reward-location associations, achieving a hit rate of 87% with robust memory discrimination (d’ = 2.49). Because d’ – a signal detection measure quantifying the standardized difference between hit and false alarm rates – provides a bias-free index of memory sensitivity, these results indicated a strong ability to distinguish reward from no reward locations. Interestingly, memory performance varied significantly across pillar locations, following a cubic rather than a linear pattern. Memory discrimination (d’) was highest for the central pillar location (P3), significantly exceeding performance for all other positions. In contrast, P1 showed the lowest d’ and the highest response bias (β), suggesting participants were more conservative in identifying P1 as the target location. This heightened bias may reflect greater difficulty discriminating P1 from other pillar locations, potentially due to its visual similarity to other pillars and its position at the track entrance where spatial distinctiveness might be reduced. P3 was also identified faster than other locations during both correct and incorrect responses, further supporting greater memory encoding and retrieval performance for this central position. It is worth noting that these findings contrast with many existing memory studies using non-spatial serial memorization list tasks, which report superior performance (and elevated ERP amplitudes) for the first and last items in a list (i.e., primacy and recency effects; Galli et al., 2012; Sederberg et al., 2006; Wiswede et al., 2007). The observed advantage for the middle positions along the track likely reflects the visuo-spatial nature of the LTM task, where participants could leverage distinctive spatial features such as pillar color, texture, or relative position to enhance memory performance.

Regarding the electrophysiological data, feedback cues presented during the encoding phase elicited robust increases in RPT power (5-8 Hz) that peaked approximately 258 ms after cue onset, with bilateral posterior distribution and right hemisphere dominance, consistent with our previous work (Baker & Holroyd, 2009, 2013; Güth et al., 2025). Critically, RPT exhibited sensitivity to both cue valence and spatial location. In particular, reward cues that participants needed to memorize elicited significantly larger RPT than no reward cues, suggesting RPT tracks motivationally relevant information for memory encoding. Further, RPT showed a spatial gradient, with the earliest pillar location (P1) eliciting stronger RPT than the last location (P5). This spatial sensitivity aligns with iEEG findings showing that theta oscillations in the parahippocampal region are modulated by goal-relevant spatial information during navigation (Qasim et al., 2019; Stangl et al., 2021). During the recall stage, pillars previously associated with reward cues similarly evoked greater RPT than no reward pillars, indicating that RPT continued to distinguish motivationally salient locations during memory retrieval. The spatial gradient observed during encoding (P1 > P5) was also present during recall, suggesting stable spatial representations across encoding and recall stages. Critically, encoding-related RPT predicted subsequent memory performance. Participants with stronger RPT during reward cue encoding showed both larger d’ values (better discrimination between target and non-target locations) and higher β values (more conservative responding, fewer false alarms). A significant interaction between d’ and β in predicting RPT power revealed a synergistic relationship: Increases in both memory measures were associated with the highest RPT power. This pattern indicates that stronger RPT during encoding supported more accurate and more cautious memory-guided decisions during retrieval. Taken together, these observations support the hypothesis that RPT reflects memory encoding and recall.

Interestingly, while RPT during encoding predicted memory outcomes, individual differences in RPT during the recall stage did not predict memory performance. This contrasts with Qasim et al. (2019), who found that entorhinal cortex firing during recall trials selectively tracked cued memories and predicted successful retrieval. A key methodological difference may account for this discrepancy: in Qasim’s task, participants navigated the virtual track during recall while objects were invisible, requiring them to engage spatial memory systems while moving through the previously encoded environment. In contrast, our recall stage presented static images of pillar locations without navigation. Without active movement through the spatial context, recall may not engage parahippocampal theta mechanisms to the same degree. This may explain why recall-related RPT did not predict individual memory performance, despite the clear predictive power of encoding-related RPT. It is also worth noting that the spatial pattern of RPT power dissociated from memory performance. While RPT decreased from P1 to P5, memory performance followed a cubic pattern with peak performance at the central location (P3). This dissociation suggests that RPT could reflect early encoding processes modulated by spatial position along the track, which is potentially linked to an attentional orienting response. In addition, the RPT power decrease could be due to repeated stimulus presentation at short inter-stimulus intervals, resulting in a decline in phase-locked evoked potentials after the first stimulus (supplementary Figure S5), consistent with neural adaptations observed in serial stimulus EEG paradigms (Agam & Sekuler, 2007; Schweinberger & Neumann, 2016; Summerfield et al., 2011). By contrast, actual memory formation might additionally depend on distinctive spatial features. For example, the central pillar location may have benefited from richer contextual cues, as it was situated between other pillar locations along the track, providing more distinctive visual features. This interpretation aligns with computational models proposing that successful spatial memory relies not only on theta phase coding of individual locations but also on the integration of multiple spatial cues and contextual information (Burgess, 2008; Hasselmo, 2008).

## Limitations

The results of the current study support the hypothesis that RPT is related to spatial memory functions and that elevated RPT power can differentiate between encoding of information relevant for goal attainment (reward cues) and nonrelevant information (no reward cues). Yet, this latter finding might be affected by the difference in stimulus probability between reward and no reward cues. The frequency of an event in EEG paradigms is a well-known influence on the amplitude of EEG signals, as documented for example in oddball paradigms (Courchesne et al., 1975; Sutton et al., 1965). Such oddball effects are primarily reported for late components over central and posterior electrodes such as the P3a and P3b (Polich, 2007; Squires et al., 1975). Still, some evidence suggests that earlier lateralized components such as the N170 and the visual mismatch negativity during face processing are affected by stimulus probability as well (Astikainen & Hietanen, 2009; Li et al., 2012; Liu et al., 2025; Susac et al., 2004; Zhao & Li, 2006). Therefore, the valence difference in the encoding stage when reward cues are presented once per trial and no reward cues four times per trial might be influenced by stimulus probability. For future research, this task could be amended by increasing the number of reward cues per trial. Additionally, increasing the number of reward cues could offer another tool to vary the difficulty of the LTM task. Allowing a variable number of reward cues per trial increases memory load by requiring more positions to be encoded and creating greater uncertainty about which pillars contain reward cues. With the current design, remembering the location of the reward cue is sufficient to exclude all other pillars on the track.

An important unresolved question concerns the role of beta power increases (13-30 Hz) that coincided with RPT during the recall stage. Prefrontal and posterior theta power as well as cross-frequency coupling between theta and low gamma oscillations (30-60 Hz) in human EEG have long been discussed as a correlate of memory retrieval (Belluscio et al., 2012; Canolty et al., 2006; Jacobs et al., 2006; Kaplan et al., 2014; Mormann et al., 2005; Nyhus & Curran, 2010; Sederberg et al., 2003). While beta power has received less attention than low gamma oscillations, beta has been mentioned alongside theta and gamma power for predicting memory recall (Jacobs et al., 2006; Nyhus, 2018; Sederberg et al., 2003). Specifically, the simultaneous occurrence of RPT and beta activity when participants were presented with the pillars from the encoding stage, could be indicative of postretrieval processes such as alignment of behavioral responses with task goals (Nyhus, 2018). An analysis and more detailed discussion of cross-frequency couplings during the recall stage are provided in the supplementary materials. However, since this analysis was an ad hoc exploration, the functional significance, frequency specificity, temporal dynamics, and spatial distribution of these couplings in the LTM task remain to be determined and merit investigation in future research.

## Conclusions

The current study establishes RPT as a viable scalp-recorded marker of human spatial memory encoding and retrieval during navigation. Across multiple replications, RPT has proven to be a robust signal associated with spatial memory and linked to medial temporal activity, specifically in the PHG (Baker & Holroyd, 2013; Güth et al., 2025). These findings provide an important bridge between animal models, human iEEG recordings, and non-invasive EEG research on spatial memory systems. The ability to track medial temporal lobe memory signals non-invasively on the scalp offers a promising tool for clinical applications in both the diagnosis and treatment of memory-related disorders. Such markers could enhance early detection and personalized treatment approaches by providing objective, physiological indicators of memory function. Future work should examine whether RPT is sensitive to memory-related disorders such as Alzheimer’s disease, or to pharmacological and transcranial magnetic stimulation interventions, to evaluate its potential as a biomarker of disorder onset, progression, and treatment response.

## Acknowledgements

We thank G. Karpov, N. Lalta, M.-H. Lin, and J. Stringfellow for assistance with the data collection.

## Author contributions

Malte R. Güth: conceptualization, software, investigation, formal analysis, visualization, writing – original draft, writing – review & editing. Travis E. Baker: conceptualization, methodology, software, supervision, writing – review & editing.

## References

Agam, Y., & Sekuler, R. (2007). Interactions between working memory and visual perception: An ERP/EEG study. NeuroImage, 36(3), 933–942. 10.1016/j.neuroimage.2007.04.014

Aghajan, Z. M., Schuette, P., Fields, T. A., Tran, M. E., Siddiqui, S. M., Hasulak, N. R., Tcheng, T. K., Eliashiv, D., Mankin, E. A., Stern, J., Fried, I., & Suthana, N. (2017). Theta Oscillations in the Human Medial Temporal Lobe during Real-World Ambulatory Movement. Current Biology, 27(24), Article 24. 10.1016/j.cub.2017.10.062

Astikainen, P., & Hietanen, J. K. (2009). Event-related potentials to task-irrelevant changes in facial expressions. Behavioral and Brain Functions, 5(1), 30. 10.1186/1744-9081-5-30

Baker, T. E., & Holroyd, C. B. (2009). Which Way Do I Go? Neural Activation in Response to Feedback and Spatial Processing in a Virtual T-Maze. Cerebral Cortex, 19(8), Article 8. 10.1093/cercor/bhn223

Baker, T. E., & Holroyd, C. B. (2013). The topographical N170: Electrophysiological evidence of a neural mechanism for human spatial navigation. Biological Psychology, 94(1), Article 1. 10.1016/J.BIOPSYCHO.2013.05.004

Baker, T. E., Umemoto, A., Krawitz, A., & Holroyd, C. B. (2015). Rightward-biased hemodynamic response of the parahippocampal system during virtual navigation. Scientific Reports, 5(1), Article 1. 10.1038/srep09063

Belluscio, M. A., Mizuseki, K., Schmidt, R., Kempter, R., & Buzsaki, G. (2012). Cross-Frequency Phase-Phase Coupling between Theta and Gamma Oscillations in the Hippocampus. Journal of Neuroscience, 32(2), Article 2. 10.1523/JNEUROSCI.4122-11.2012

Burgess, N. (2008). Grid cells and theta as oscillatory interference: Theory and predictions. Hippocampus, 18(12), Article 12. 10.1002/hipo.20518

Buzsáki, G. (2005). Theta rhythm of navigation: Link between path integration and landmark navigation, episodic and semantic memory. Hippocampus, 15(7), Article 7. 10.1002/hipo.20113

Buzsáki, G., & Moser, E. I. (2013). Memory, navigation and theta rhythm in the hippocampal-entorhinal system. Nature Neuroscience, 16(2), Article 2. 10.1038/nn.3304

Canolty, R., Edwards, E., Dalal, S., & Soltani, M. (2006). High Gamma Power Is Phase-Locked to Theta Oscillations in Human Neocortex. Science. https://www.researchgate.net/profile/Srikantan_Nagarajan/publication/6817257_High_gamma_power_is_phase-locked_to_theta_oscillations_in_human_neocortex/links/00b4953973ad6c9ee2000000.pdf

Courchesne, E., Hillyard, S. A., & Galambos, R. (1975). Stimulus novelty, task relevance and the visual evoked potential in man. Electroencephalography and Clinical Neurophysiology, 39(2), 131–143. 10.1016/0013-4694(75)90003-6

Ekstrom, A. D., Caplan, J. B., Ho, E., Shattuck, K., Fried, I., & Kahana, M. J. (2005). Human hippocampal theta activity during virtual navigation. Hippocampus, 15(7), Article 7. 10.1002/hipo.20109

Galli, G., Leng Choy, T., & Otten, L. J. (2012). Prestimulus brain activity predicts primacy in list learning. Cognitive Neuroscience, 3(3–4), 160–167. 10.1080/17588928.2012.670105

Givens, B. (1996). Stimulus-evoked resetting of the dentate theta rhythm: Relation to working memory. NeuroReport, 8(1), Article 1.

Goyal, A., Miller, J., Qasim, S. E., Watrous, A. J., Zhang, H., Stein, J. M., Inman, C. S., Gross, R. E., Willie, J. T., & Lega, B. (2020). Functionally distinct high and low theta oscillations in the human hippocampus. Nature Communications, 11(1), Article 1.

Gramfort, A., Schurger, A., & Naccache, L. (2014). Two Distinct Dynamic Modes Subtend the Detection of Unexpected Sounds. 9(1). 10.1371/journal.pone.0085791

Green, D. M., & Swets, J. A. (1974). Signal detection theory and psychophysics (pp. xiii, 479).

Robert E. Krieger. Greenhouse, S. W., & Geisser, S. (1959). On methods in the analysis of profile data. Psychometrika, 24(2), Article 2. 10.1007/BF02289823

Güth, M. R., Reid, A., Zhang, Y., Huntgeburth, S. C., Mill, R. D., Dagher, A., Kerns, K., Holroyd, C. B., Petrides, M., Cole, M. W., & Baker, T. E. (2025). Right posterior theta reflects human parahippocampal phase resetting by salient cues during goal-directed navigation. Imaging Neuroscience. 10.1162/IMAG.a.105

Hasselmo, M. E. (2008). Grid cell mechanisms and function: Contributions of entorhinal persistent spiking and phase resetting. Hippocampus, 18(12), Article 12. 10.1002/hipo.20512

Hasselmo, M. E., & Stern, C. E. (2014). Theta rhythm and the encoding and retrieval of space and time. NeuroImage, 85 Pt 2, 656–666. 10.1016/j.neuroimage.2013.06.022

Hegarty, M., Richardson, A. E., Montello, D. R., Lovelace, K., & Subbiah, I. (2002). Development of a self-report measure of environmental spatial ability. Intelligence, 30(5), Article 5. 10.1016/S0160-2896(02)00116-2

Jacobs, J., Hwang, G., Curran, T., & Kahana, M. J. (2006). EEG oscillations and recognition memory: Theta correlates of memory retrieval and decision making. NeuroImage, 32(2), Article 2. 10.1016/j.neuroimage.2006.02.018

Jacobs, J., Kahana, M. J., Ekstrom, A. D., Mollison, M. V., & Fried, I. (2010). A sense of direction in human entorhinal cortex. Proceedings of the National Academy of Sciences of the United States of America, 107(14), Article 14. 10.1073/pnas.0911213107

Jacobs, J., Weidemann, C. T., Miller, J. F., Solway, A., Burke, J. F., Wei, X.-X., Suthana, N., Sperling, M. R., Sharan, A. D., Fried, I., & Kahana, M. J. (2013). Direct recordings of grid-like neuronal activity in human spatial navigation. Nature Neuroscience, 16(9), Article 9. 10.1038/nn.3466

Jasper, H. H. (1958). The 10-20 electrode system of the International Federation. Electroencephalogr Clin Neurophysiol, 10, 370–375.

Kaplan, R., Bush, D., Bonnefond, M., Bandettini, P. A., Barnes, G. R., Doeller, C. F., & Burgess, N. (2014). Medial prefrontal theta phase coupling during spatial memory retrieval. Hippocampus, 24(6), Article 6. 10.1002/hipo.22255

Li, X., Lu, Y., Sun, G., Gao, L., & Zhao, L. (2012). Visual mismatch negativity elicited by facial expressions: New evidence from the equiprobable paradigm. Behavioral and Brain Functions, 8(1), 7. 10.1186/1744-9081-8-7

Liu, X., Li, X., & Astikainen, P. (2025). N170 Amplitude to Rare Neutral Faces in an Oddball Condition Reflects Prediction Error. The European Journal of Neuroscience, 62(7), e70264. 10.1111/ejn.70264

Macmillan, N. A., & Creelman, C. D. (1990). Response bias: Characteristics of detection theory, threshold theory, and “non-parametric” indexes. Psychological Bulletin, 401–413.

McCartney, H., Johnson, A. D., Weil, Z. M., & Givens, B. (2004). Theta reset produces optimal conditions for long-term potentiation. Hippocampus, 14(6), Article 6.

Mormann, F., Fell, J., Axmacher, N., Weber, B., Lehnertz, K., Elger, C. E., & Fernández, G. (2005). Phase/amplitude reset and theta–gamma interaction in the human medial temporal lobe during a continuous word recognition memory task. Hippocampus, 15(7), Article 7.

Nadasdy, Z., Nguyen, T. P., Török, Á., Shen, J. Y., Briggs, D. E., Modur, P. N., & Buchanan, R. J. (2017). Context-dependent spatially periodic activity in the human entorhinal cortex.Proceedings of the National Academy of Sciences, 114(17), Article 17. 10.1073/pnas.1701352114

Nyhus, E. (2018). Brain Networks Related to Beta Oscillatory Activity during Episodic Memory Retrieval. Journal of Cognitive Neuroscience, 30(2), 174–187. 10.1162/jocn_a_01194

Nyhus, E., & Curran, T. (2010). Functional role of gamma and theta oscillations in episodic memory. *Neuroscience & Biobehavioral Reviews*, Binding Processes: Neurodynamics and Functional Role in Memory and Action, 34(7), 1023–1035. 10.1016/j.neubiorev.2009.12.014

Oldfield, R. C. (1971). The assessment and analysis of handedness: The Edinburgh inventory. Neuropsychologia, 9(1), Article 1. 10.1016/0028-3932(71)90067-4

Polich, J. (2007). Updating P300: An integrative theory of P3a and P3b. Clinical Neurophysiology. http://www.sciencedirect.com/science/article/pii/S1388245707001897

Qasim, S. E., Miller, J., Inman, C. S., Gross, R. E., Willie, J. T., Lega, B., Lin, J.-J., Sharan, A., Wu, C., Sperling, M. R., Sheth, S. A., McKhann, G. M., Smith, E. H., Schevon, C., Stein, J. M., & Jacobs, J. (2019). Memory retrieval modulates spatial tuning of single neurons in the human entorhinal cortex. Nature Neuroscience, 22(12), Article 12. 10.1038/s41593-019-0523-z

R Development Core Team. (2016). R: A language and environment for statistical computing. R Foundation for Statistical Computing, Vienna, Austria. 2014.

Rizzuto, D. S., Madsen, J. R., Bromfield, E. B., Schulze-Bonhage, A., Seelig, D., Aschenbrenner-Scheibe, R., & Kahana, M. J. (2003). Reset of human neocortical oscillations during a working memory task. Proceedings of the National Academy of Sciences, 100(13), 7931–7936. 10.1073/pnas.0732061100

Schweinberger, S. R., & Neumann, M. F. (2016). Repetition effects in human ERPs to faces. *Cortex*, Special Issue:Repetition Suppression-an Integrative View, 80, 141–153. 10.1016/j.cortex.2015.11.001

Seabold, S., & Perktold, J. (2010). Statsmodels: Econometric and statistical modeling with python. Proceedings of the 9th Python in Science Conference, 57, 61.

Sederberg, P. B., Gauthier, L. V., Terushkin, V., Miller, J. F., Barnathan, J. A., & Kahana, M. J. (2006). Oscillatory correlates of the primacy effect in episodic memory. NeuroImage, 32(3), 1422–1431. 10.1016/j.neuroimage.2006.04.223

Sederberg, P. B., Kahana, M. J., Howard, M. W., Donner, E. J., & Madsen, J. R. (2003). Theta and Gamma Oscillations during Encoding Predict Subsequent Recall. The Journal of Neuroscience, 23(34), 10809–10814. 10.1523/JNEUROSCI.23-34-10809.2003

Squires, N. K., Squires, K. C., & Hillyard, S. A. (1975). Two varieties of long-latency positive waves evoked by unpredictable auditory stimuli in man. Electroencephalography and Clinical Neurophysiology, 38(4), 387–401. 10.1016/0013-4694(75)90263-1

Stangl, M., Topalovic, U., Inman, C. S., Hiller, S., Villaroman, D., Aghajan, Z. M., Christov-Moore, L., Hasulak, N. R., Rao, V. R., Halpern, C. H., Eliashiv, D., Fried, I., & Suthana, N. (2021). Boundary-anchored neural mechanisms of location-encoding for self and others. Nature, 589(7842), Article 7842. 10.1038/s41586-020-03073-y

Summerfield, C., Wyart, V., Mareike Johnen, V., & de Gardelle, V. (2011). Human Scalp Electroencephalography Reveals that Repetition Suppression Varies with Expectation. Frontiers in Human Neuroscience, 5. 10.3389/fnhum.2011.00067

Susac, A., Ilmoniemi, R. J., Pihko, E., & Supek, S. (2004). Neurodynamic Studies on Emotional and Inverted Faces in an Oddball Paradigm. Brain Topography, 16(4), 265–268. 10.1023/B:BRAT.0000032863.39907.cb

Sutton, S., Braren, M., Zubin, J., & John, E. R. (1965). Evoked-Potential Correlates of Stimulus Uncertainty. Science, 150(3700), 1187–1188. 10.1126/science.150.3700.1187

Vallat, R. (2018). Pingouin: Statistics in Python. Journal of Open Source Software, 3(31), Article 31. 10.21105/joss.01026

Virtanen, P., Gommers, R., Oliphant, T. E., Haberland, M., Reddy, T., Cournapeau, D., Burovski, E., Peterson, P., Weckesser, W., Bright, J., Van Der Walt, S. J., Brett, M., Wilson, J., Millman, K. J., Mayorov, N., Nelson, A. R. J., Jones, E., Kern, R., Larson, E., … Vázquez-Baeza, Y. (2020). SciPy 1.0: Fundamental algorithms for scientific computing in Python. Nature Methods, 17(3), 261–272. 10.1038/s41592-019-0686-2

Williams, J. M., & Givens, B. (2003). Stimulation-induced reset of hippocampal theta in the freely performing rat. Hippocampus, 13(1), Article 1.

Winkler, I., Debener, S., Müller, K., & Tangermann, M. (2015). On the influence of high-pass filtering on ICA-based artifact reduction in EEG-ERP. 2015 37th Annual International Conference of the IEEE Engineering in Medicine and Biology Society *(EMBC)*, 4101–4105. 10.1109/EMBC.2015.7319296

Wiswede, D., Rüsseler, J., & Münte, T. F. (2007). Serial position effects in free memory recall—An ERP-study. Biological Psychology, 75(2), 185–193. 10.1016/j.biopsycho.2007.02.002

Zhao, L., & Li, J. (2006). Visual mismatch negativity elicited by facial expressions under non-attentional condition. Neuroscience Letters, 410(2), 126–131. 10.1016/j.neulet.2006.09.081

