## Supplementary materials for "Right posterior theta facilitates memory encoding and recall during virtual navigation"

#### Response bias across pillar locations

Using the same one-way repeated-measures ANOVA as for d', with Pillar Location as a within-subject factor, we observed a significant main effect on β scores, $F(4, 104) = 12.1$, $p = 4.107 \times 10^{-8}$, $\eta_p^2 = 0.32$. The first pillar P1 (M = 16.19, SD = 16.43) had significantly higher β values than all remaining pillar locations (P2: M = 4.86, SD = 2.78; $t(26) = 3.61$, $p = .012$, $d = 0.71$; P3: M = 6.62, SD = 4.1; $t(26) = 3.34$, $p = .018$, $d = 0.66$; P4: M = 4.04, SD = 1.88; $t(26) = 3.74$, $p = .009$, $d = 0.73$; P5: M = 5.19, SD = 2.83; $t(26) = 3.49$, $p = .014$, $d = 0.69$, Bonferroni-Holm corrected). This finding suggests that participants were more hesitant to identify P1 as the target pillar with the reward cue. Given that participants scored lower d' values for P1 compared to other pillars, perhaps P1 was perceived to be more difficult to differentiate from the other pillars.

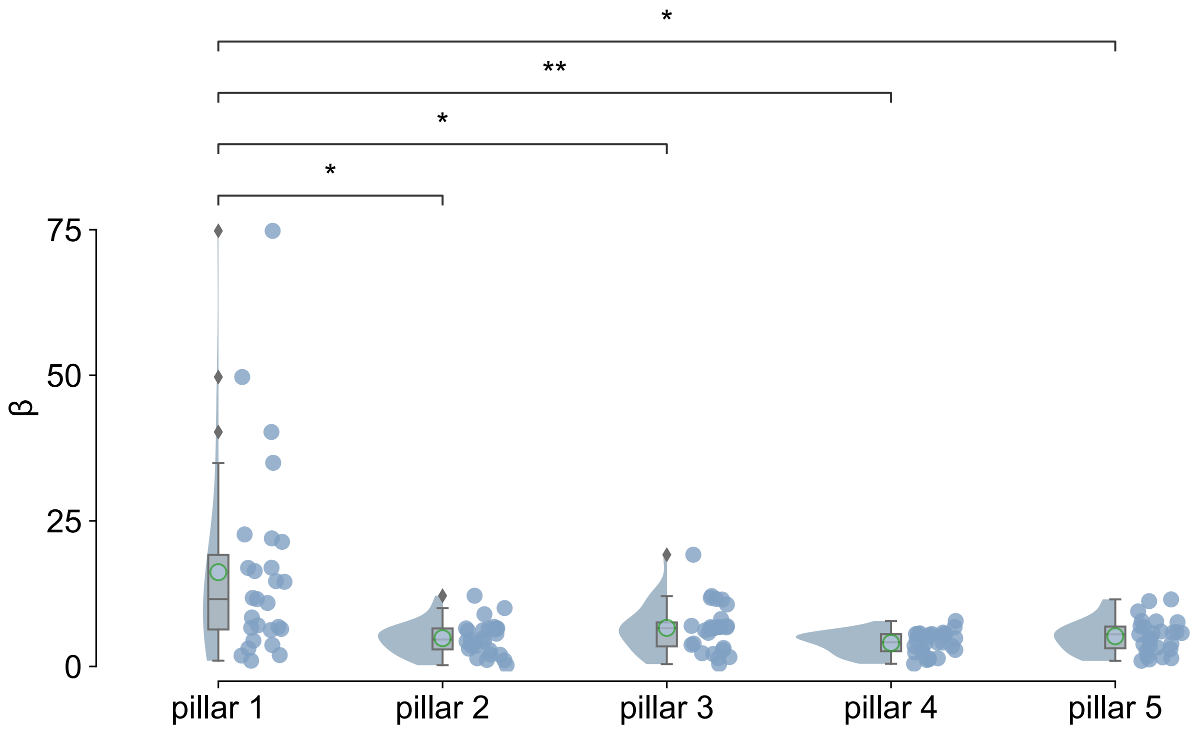


**Figure S1** Response bias (β) scores for each pillar location when the reward cue was presented at a given pillar. Raincloud plots are constructed equivalently to Figure 2. **p < .05, **p < .01*

#### Encoding-related event-related potentials

Figure S2 shows the ERP recorded over bilateral parieto-occipital channels when time-locking signals to the onset of a trial after subjects pressed the go button. When reaching each pillar location (P1-P5) and being presented with a feedback-related cue (reward, no reward), there was a bilateral negative deflection peaking after approximately 220 ms (SD = 15 ms) at P8 (M =

2.76 μV, SD = 2.77 μV) and P7 (M = 2.84 μV, SD = 2.04 μV). The negative amplitude, latency, topography, and experimental context are consistent with our previous work on the N170 during the encoding of goal-related information during spatial navigation in a virtual T-maze task (Baker & Holroyd, 2013; Güth et al., 2025).

We conducted the same three-way repeated measures ANOVA as for the RPT power data in the main manuscript with N170 amplitude as the dependent variable. N170 amplitudes were extracted from the cleaned segments as the maximum negative amplitude between 190 ms and 250 ms after the cue presentation. This ANOVA included the factors Valence (reward cue, no reward cue), Channel (P7, P8), and Pillar Location (P1, P2, P3, P4, P5). Consistent with the results for RPT power, there was a significant main effect of Valence ($F(1, 26) = 38.86$, $p = 1.35 \times 10^{-6}$, $\eta_p^2 = 0.6$). Reward cues elicited significantly larger N170 amplitudes (M = 3.62 μV, SD = 1.99 μV) compared to no reward cues (M = 1.98 μV, SD = 1.08 μV). In contrast to RPT power, there were no significant main effects for the factors Channel ($F(1, 26) = 0.03$, $p = .862$, $\eta_p^2 = 0.001$) and Pillar Location ($F(4, 104) = 1.87$, $p = .161$, $\eta_p^2 = 0.07$).

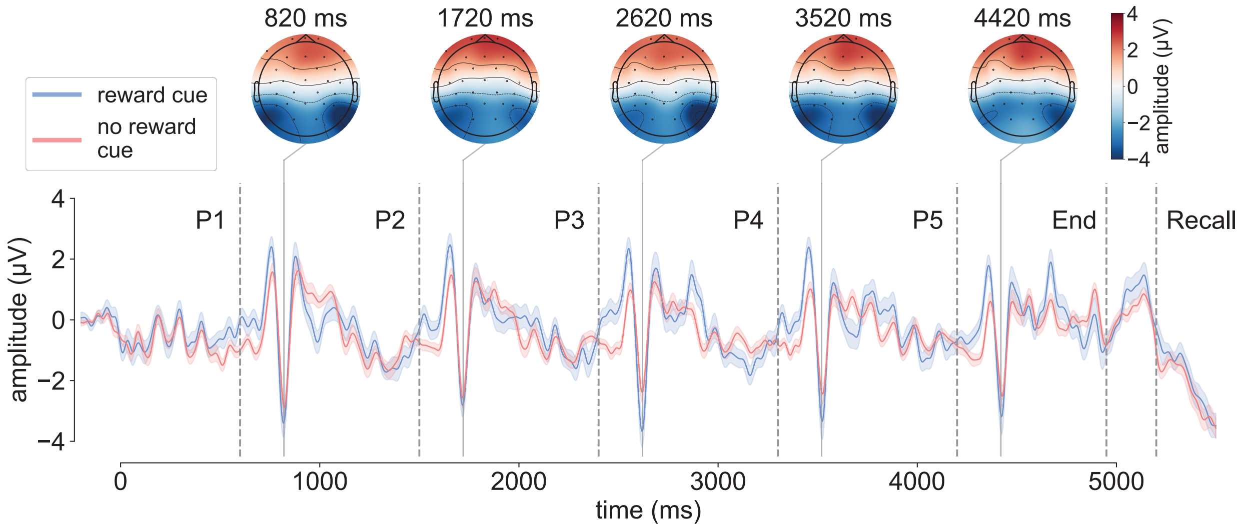


**Figure S2** Average event-related potential (ERP) at channel P8 time-locked to the beginning of a trial separated by cue valence (reward: blue, no reward: red). Identical to Figure 3, dashed lines (P1-P5) highlight the onset of cues at the five pillar pairs. End marks the arrival at the wall at the end of the linear track and the end of the encoding stage. Recall marks the onset of the recall stage. Topographies at the top depict the average amplitude values for reward cues at their respective timings noted above. Shaded areas represent ± one standard error of the mean.

**Cross-frequency coupling during recall**

As the time-frequency analysis of recall-related RPT yielded notable beta power increases (13-30 Hz) coinciding with the RPT power, we conducted an ad-hoc analysis of the beta power and its cross-frequency coupling with RPT during the recall stage. For this we considered two measures: amplitude-amplitude coupling (AAC) and phase-amplitude coupling (PAC) strength (Cohen, 2008; Jensen & Colgin, 2007; Voytek et al., 2013). Both the phase and amplitude of theta oscillations involved in memory processes traditionally have been examined in relation to their coupling with gamma oscillations (Belluscio et al., 2012; Canolty et al., 2006; Jacobs et al., 2006; Kaplan et al., 2014; Mormann et al., 2005; Nyhus & Curran, 2010; Sederberg et al., 2003). However, in the present study we observed increases in beta power during the presentation of pillars in the recall stage, which has also been implicated in memory retrieval (Jacobs et al., 2006; Nyhus, 2018; Sederberg et al., 2003). Thus, we analyzed both higher frequency bands prominent during recall in Figure 4 (beta: 13-30 Hz, gamma: 31-50 Hz) for the five potential target pillars.

First, to explore potential couplings of theta with the beta and gamma range, AAC between theta (5-8 Hz) and the two higher bands for the time window of interest (0 ms to 500 ms) was computed for each subject by applying bandpass filtering the EEG signal into frequency-specific bins (1 Hz steps). The analytic signal was computed using the Hilbert transform to extract the amplitude envelope. Amplitude envelope correlations between theta and high frequency bands were calculated across epochs for each frequency pair within the bins. Lastly, amplitude correlation values were normalized using a Fisher z-transformation and were then averaged across subjects. The resulting comodulogram in Figure S3 (left) highlighted that AAC primarily occurred between the high theta (7-8 Hz) and the beta band (Figure S4A, top). In a similar manner, PAC strength between theta phase and high frequency amplitude was estimated using the Modulation Index (Tort et al., 2010) as implemented in Tensorpac (version 0.6.5, Combrisson et al., 2020). The Hilbert transform was used to extract the instantaneous phase and amplitude. PAC values were computed for all epochs and each frequency pair for each subject within a wider time window of interest (-500 ms to 500 ms) than for the AAC analysis to obtain more stable PAC estimates given the slow theta frequencies. PAC values were averaged across subjects to obtain the comodulogram in Figure S3 (right). Low theta (5-6 Hz) was coupled to the full high frequency range.

Second, to investigate time-varying couplings in response to the pillar presentation theta-beta and theta-gamma AAC and PAC values were calculated for separate time bins (Figure S4). To this end, overlapping windows of 500 ms were defined and moved across the full epoch (±2500 ms) in steps of 100 ms. This window length provides a balance between temporal resolution and inclusion of at least 2.5 theta cycles for coupling estimates. We used the identified optimal pairings between low frequency ranges (AAC: 7-8 Hz, PAC: 5-6 Hz) and high frequency ranges ($AAC_{beta}$: 13-20 Hz, $PAC_{beta}$: 13-30 Hz; $AAC_{gamma}$: 31-50 Hz, $PAC_{gamma}$: 31-50 Hz) in Figure S3. For each subject the resulting AAC and PAC time-series were averaged across epochs.

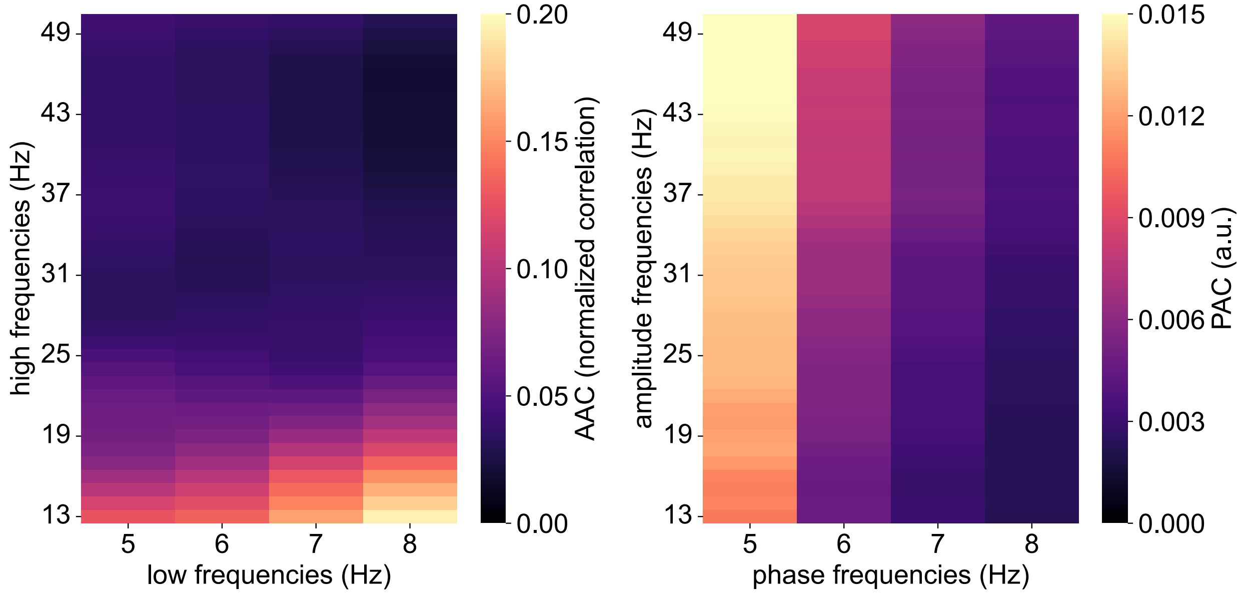


**Figure S3** Comodulograms of amplitude-amplitude coupling (AAC, left) and phase-amplitude coupling (PAC, right) between low frequencies (theta: 5-8 Hz) and high frequencies (beta: 13-30 Hz, gamma: 31-50 Hz). Color denotes the unitless coupling strength for PAC and Fisher z-transformed correlation coefficients for AAC.

Figure S4A shows the time courses of theta-beta and theta-gamma AAC and PAC values aligned to the pillar presentations during the recall stage. Theta-beta AAC and PAC peaked within the first 500 ms following the pillar presentation while theta-gamma couplings did not show clear event-related changes. To test the relationship between cross-frequency coupling and memory measures peak AAC and PAC values within the first 500 ms after pillar presentation were extracted for each subject and entered as regressors into GLMs to predict d’ and β values (Figure S4B). Theta-beta AAC did not significantly predict d’ ($t(26) = -1.02$, $p = .308$, $b = -0.256$, $CI_{95\%} = -0.747$-0.236), but it did significantly predict β with a negative regression coefficient ($t(26) = -1.99$, $p = .047$, $b = -0.43$, $CI_{95\%} = -0.854$-0.005), indicating a more liberal response bias (less cautious target pillar identification) in subjects with stronger theta-beta coupling. To examine whether theta-beta AAC explains variance in β beyond that accounted for by encoding-related RPT power, we fitted an additional GLM including both predictors. By entering both theta-beta AAC and RPT power, the model estimates the unique contribution of each predictor while controlling for their shared variance. In this model, neither predictor significantly predicted β, indicating largely shared variance between encoding-related RPT and theta-beta AAC regressors (RPT: $t(26) = 0.81$, $p = .418$, $b = 0.181$, $CI_{95\%} = -0.257$-$0.62$; theta-beta AAC: $t(26) = 1.79$, $p = .074$, $b = -0.397$, $CI_{95\%} = -0.832$-$0.039$).

Regarding theta-gamma couplings, AAC significantly predicted d’ (Figure S4B, top left; $t(26) = 2.33$, $p = .02$, $b = 0.53$, $CI_{95\%} = 0.085$-$0.975$) but not β ($t(26) = 1.21$, $p = .228$, $b = 0.276$, $CI_{95\%} = -0.173$-$0.726$). Equivalently to the analysis above, theta-gamma AAC and encoding RPT power were entered into another GLM together to test if AAC significantly predicts d’ while controlling for the shared variance with RPT power. Unlike for theta-beta AAC, the effect of theta-gamma both AAC ($t(26) = 2.94$, $p = .003$, $b = 0.563$, $CI_{95\%} = 0.187$-$0.94$) and the RPT predictor ($t(26) = 2.95$, $p = .003$, $b = 0.571$, $CI_{95\%} = 0.192$-$0.949$) remained significant. This indicates that each of their contributions were unique. No significant relationships between PAC values and memory measures were found (d’: $t(26) = 0.02$, $p = .988$, $b = 0.004$, $CI_{95\%} = -0.501$-$0.509$; β: $t(26) = -0.1$, $p = .924$, $b = -0.023$, $CI_{95\%} = -0.489$-$0.443$).

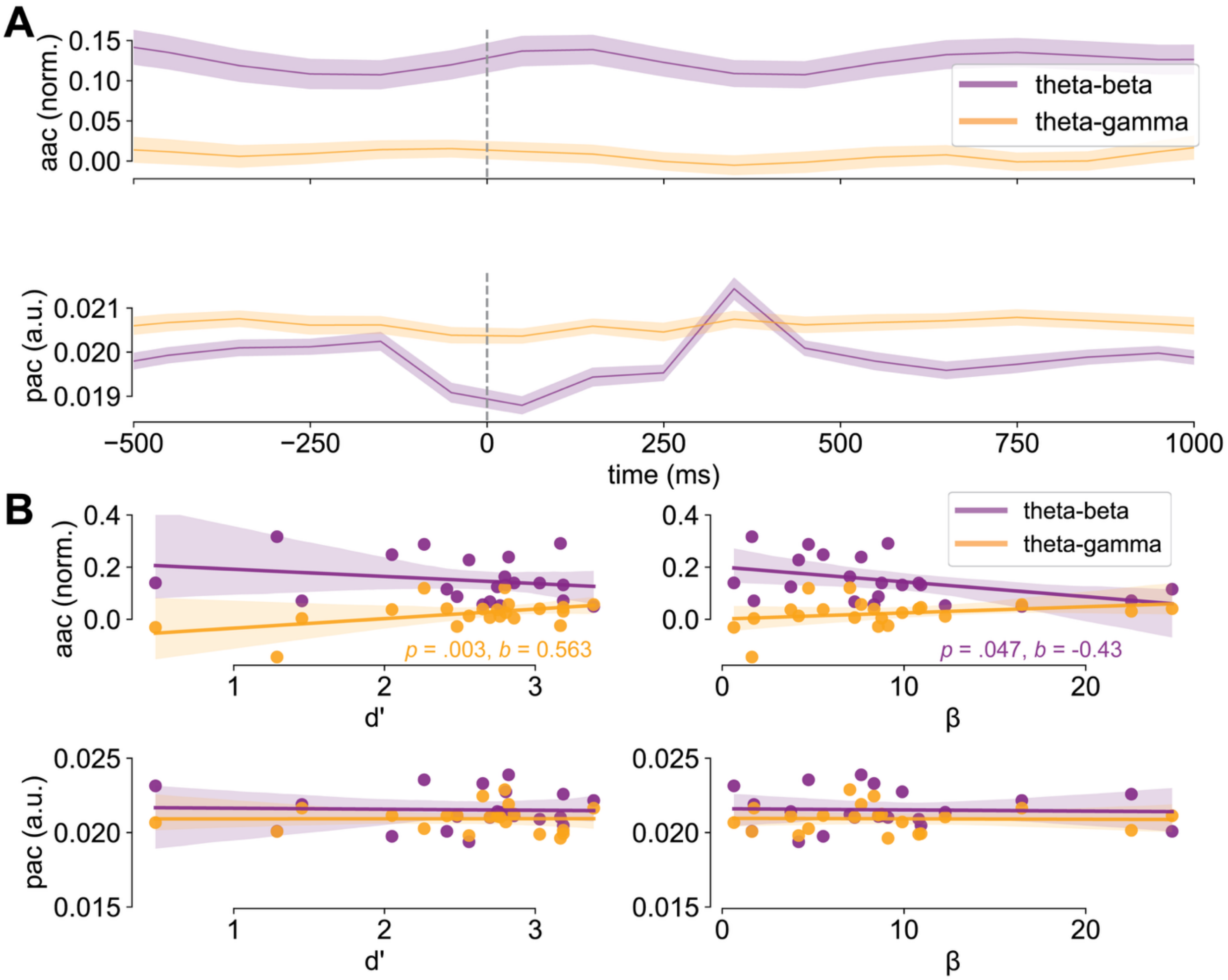


**Figure S4** **A**: Time-courses of theta-beta AAC (top, normalized correlations) and PAC (bottom) values time-locked to the pillar presentation (dashed line) averaged across subjects. Values are plotted separately for theta-beta (magenta) and theta-gamma (orange) couplings. Shaded areas mark ± one standard error of the mean. **B**: Regression plots for the relationship of theta-beta AAC (top row) and PAC (bottom row) with memory measures, d' (left column) and β scores (right column). Colors denote theta-beta and theta-gamma values like in **A**. Shaded areas are a 95% confidence interval around the regression line. Significant coefficients and p-values are color-coded for beta and gamma couplings.

Taken together, these AAC and PAC dynamics reveal small event-related changes in cross-frequency coupling when participants engaged in memory retrieval. Although PAC did not have significant relationships with memory performance, theta-beta AAC predicted response bias with a negative coefficient and theta-gamma AAC predicted d' with a positive coefficient. Regarding theta-beta AAC, participants with high coupling were more likely to identify pillars as the target pillar, risking more false positives. Regarding theta-gamma AAC, participants with high coupling scored higher discriminability values, indicating better ability to differentiate between the target pillar and others. Notably, only the relationship between theta-gamma AAC and d' was significant when controlling for the shared variance with RPT during encoding. This entails that variance in memory performance explained by theta-beta AAC is not unique and largely overlaps with variability captured by encoding RPT. Thus, enhanced RPT during encoding and reduced theta-beta AAC during memory retrieval may reflect related processes.

Theta-gamma AAC appears to support memory recall and discrimination performance beyond the contribution of encoding RPT, potentially by coordinating low and high frequency activity across the medial temporal lobe. While memory-related theta amplitude, as indexed by RPT, increases during recall (Vivekananda et al., 2021), gamma amplitude increases have been linked to the processing of object-location information (Neves et al., 2022), as would be required for the recall stage in the LTM. Consistent with this interpretation, specifically hippocampal theta-gamma PAC is commonly observed in humans and rodents during memory engagement, particularly during recall (Mormann et al., 2005; Tamura et al., 2017). More generally, the comodulation of theta and gamma amplitudes may facilitate the coordination of neural processes necessary for the successful recall of goal-relevant information tied to specific spatial contexts such as pillar locations (Shirvalkar et al., 2010). In rodents, increased concomitant theta and gamma activity during memory tasks has been associated with improved discrimination performance (Neves et al., 2022). Hippocampal theta and gamma oscillatory interactions during exploration of displaced compared to stationary objects predicted successful discrimination performance in rats.

This pattern of results suggests an inverse relationship between theta-beta AAC and RPT. While increased RPT predicted greater response bias, reflecting increased caution or perhaps certainty about the target pillar, stronger theta-beta AAC was associated with reduced bias and less careful responding. This aligns with simultaneous EEG-fMRI evidence showing that increased posterior beta power during memory retrieval correlates with novelty in word memorization task and reduced posterior parietal BOLD activation (Nyhus, 2018). In that study, novel words elicited greater beta power than previously memorized words, which was interpreted as reflecting postretrieval processes involved in aligning responses with task goals. In the context of this evidence, the present theta-beta AAC may similarly relate to greater readiness to press the target response button, potentially supporting postretrieval motor response preparation. By contrast, theta-gamma AAC appears complimentary to RPT in facilitating successful memory recall for discrimination.

**Evoked RPT power decline across the track**

As described in the main manuscript, there was a significant decline in total RPT power from the first to the last location along the track regardless of EEG channel (P7, P8) or cue valence (reward, no reward). This linear decline contrasts with the cubic relationship observed between memory performance and pillar location. This power decline from the first to the last stimulus presentation resembles findings from serial presentation paradigms, in which evoked amplitudes decrease with repeated stimuli presented at fixed, short intervals (Agam & Sekuler, 2007; Schweinberger & Neumann, 2016; Summerfield et al., 2011). The observed RPT power decline may be driven by the evoked (i.e., phase-locked to the stimulus presentation) and reflect short-term adaptation to predictable repeated stimulus presentation. This possibility would align with the interpretation that RPT reflects an early attentional orienting response during memory encoding that diminishes with repetition.

To test whether a decline in evoked power was underlying the decline in total RPT we separated evoked and induced (non-phase-locked responses) spectra by subtracting the evoked RPT power from the total RPT power (for more details, see Baker & Holroyd (2013), Hajihosseini & Holroyd (2013), and Marco-Pallarés et al. (2007)). The evoked power was determined by convolving the averaged ERP with the same wavelet analysis as described in the main manuscript. The baseline correction was also performed analogously. Post-stimulus power was divided by the average of the -200 to -100 ms pre-stimulus baseline window and then the baseline average power was subtracted from the post-stimulus values. Each pillar location was corrected with its own baseline window. Figure S5 shows the time-varying power at channel P8 time-locked to the start of the track separately for evoked, induced, and total spectra. A one-way repeated measures ANOVA with the factor Pillar Location (P1, P2, P3, P4, P5) and evoked RPT power as the dependent variable yielded a significant main effect ($F(4, 108) = 4.13$, $p = .004$, $\eta_p^2 = 0.13$). Pos hoc paired t-tests revealed that P1 (M = 0.44, SD = 0.33) elicited significantly larger evoked RPT power increases than P2 (M = 0.26, SD = 0.17; $t(26) = 3.51$, $p = .016$, $d = 0.66$, Bonferroni-holm corrected), P4 (M = 0.31, SD = 0.26; $t(26) = 3.3$, $p = .025$, $d = 0.62$), and P5 (M = 0.24, SD = 0.39;

$t(26) = 3.19$, $p = .028$, $d = 0.6$). When using the induced power as the dependent variable, Pillar Location did not have a significant main effect ($F(4, 108) = 2.21$, $p = .072$, $\eta_p^2 = 0.08$).

These results indicate that only phase-locked (evoked) power declined after the first pillar, relative to the second, third, and fourth pillars. No comparable spatial modulation was observed for induced power. Thus, the reduction in total power reported in the main manuscript is likely driven by a decrease in phase-locked responses at the pillars following the first. This pattern is consistent with the interpretation of repeated stimulus presentation, leading to an adaptation of evoked activity and a diminishing orienting response during spatial memory encoding.

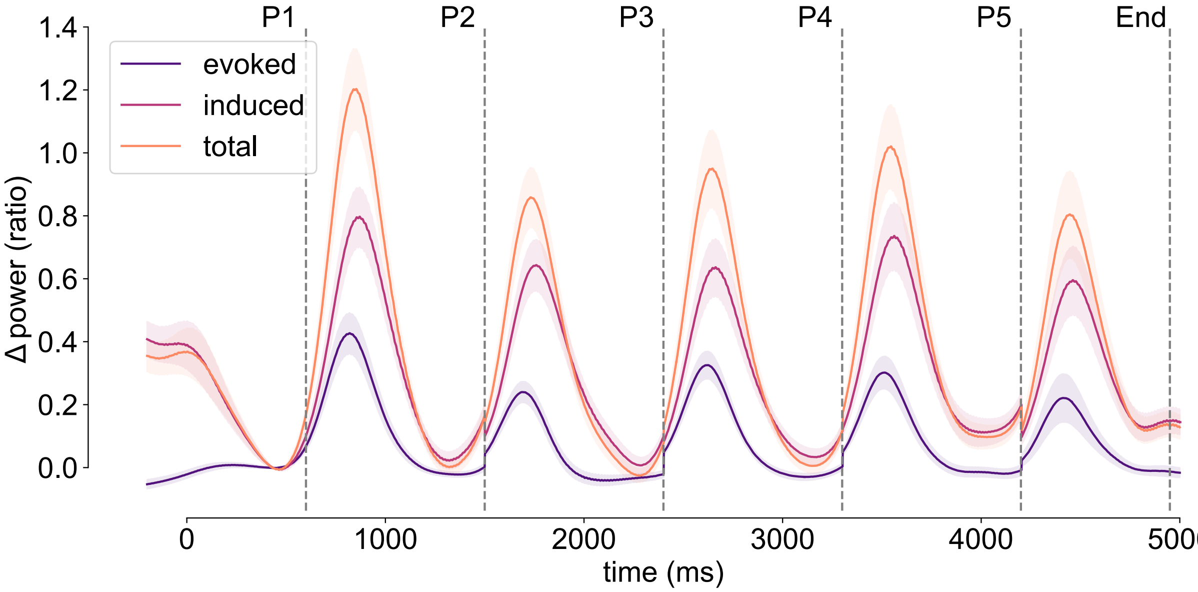


**Figure S5** Average time-courses of total, evoked, and induced power (color-coded). Dotted lines mark the timings of the cue presentation at the five pillar locations (P1, P2, P3, P4, P5) and the end of the encoding trial (End). Shaded areas mark ± one standard error of the mean.

### References

Agam, Y., & Sekuler, R. (2007). Interactions between working memory and visual perception: An ERP/EEG study. *NeuroImage*, *36*(3), 933–942. https://doi.org/10.1016/j.neuroimage.2007.04.014

Baker, T. E., & Holroyd, C. B. (2013). The topographical N170: Electrophysiological evidence of a neural mechanism for human spatial navigation. *Biological Psychology*, *94*(1), Article 1. https://doi.org/10.1016/J.BIOPSYCHO.2013.05.004

Belluscio, M. A., Mizuseki, K., Schmidt, R., Kempter, R., & Buzsaki, G. (2012). Cross-Frequency Phase-Phase Coupling between Theta and Gamma Oscillations in the

Hippocampus. *Journal of Neuroscience*, *32*(2), Article 2. https://doi.org/10.1523/JNEUROSCI.4122-11.2012

Canolty, R., Edwards, E., Dalal, S., & Soltani, M. (2006). High Gamma Power Is Phase-Locked to Theta Oscillations in Human Neocortex. *Science*. https://www.researchgate.net/profile/Srikantan_Nagarajan/publication/6817257_High_gamma_power_is_phase-locked_to_theta_oscillations_in_human_neocortex/links/00b4953973ad6c9ee2000000.pdf

Cohen, M. X. (2008). Assessing transient cross-frequency coupling in EEG data. *Journal of Neuroscience Methods*, *168*(2), Article 2. https://doi.org/10.1016/j.jneumeth.2007.10.012

Combrisson, E., Nest, T., Brovelli, A., Ince, R. A. A., Soto, J. L. P., Guillot, A., & Jerbi, K. (2020). Tensorpac: An open-source Python toolbox for tensor-based phase-amplitude coupling measurement in electrophysiological brain signals. *PLOS Computational Biology*, *16*(10), e1008302. https://doi.org/10.1371/journal.pcbi.1008302

Güth, M. R., Reid, A., Zhang, Y., Huntgeburth, S. C., Mill, R. D., Dagher, A., Kerns, K., Holroyd, C. B., Petrides, M., Cole, M. W., & Baker, T. E. (2025). Right posterior theta reflects human parahippocampal phase resetting by salient cues during goal-directed navigation. *Imaging Neuroscience*. https://doi.org/10.1162/IMAG.a.105

Hajihosseini, A., & Holroyd, C. B. (2013). Frontal midline theta and N200 amplitude reflect complementary information about expectancy and outcome evaluation. *Psychophysiology*, *50*(6), Article 6. https://doi.org/10.1111/psyp.12040

Jacobs, J., Hwang, G., Curran, T., & Kahana, M. J. (2006). EEG oscillations and recognition memory: Theta correlates of memory retrieval and decision making. *NeuroImage*, *32*(2), Article 2. https://doi.org/10.1016/j.neuroimage.2006.02.018

Jensen, O., & Colgin, L. L. (2007). Cross-frequency coupling between neuronal oscillations. *Trends in Cognitive Sciences*, *11*(7), 267–269. https://doi.org/10.1016/j.tics.2007.05.003

Kaplan, R., Bush, D., Bonnefond, M., Bandettini, P. A., Barnes, G. R., Doeller, C. F., & Burgess, N. (2014). Medial prefrontal theta phase coupling during spatial memory retrieval. *Hippocampus*, *24*(6), Article 6. https://doi.org/10.1002/hipo.22255

Marco-Pallarés, J., Müller, S. V., & Münte, T. F. (2007). Learning by doing: An fMRI study of feedback-related brain activations. *NeuroReport*, *18*(14), Article 14. https://doi.org/10.1097/WNR.0b013e3282e9a58c

Mormann, F., Fell, J., Axmacher, N., Weber, B., Lehnertz, K., Elger, C. E., & Fernández, G. (2005). Phase/amplitude reset and theta–gamma interaction in the human medial temporal lobe during a continuous word recognition memory task. *Hippocampus*, *15*(7), Article 7.

Neves, L., Lobão-Soares, B., Araujo, A. P. de C., Furtunato, A. M. B., Paiva, I., Souza, N., Morais, A. K., Nascimento, G., Gavioli, E., Tort, A. B. L., Barbosa, F. F., & Belchior, H. (2022). Theta and gamma oscillations in the rat hippocampus support the discrimination of object displacement in a recognition memory task. *Frontiers in Behavioral Neuroscience*, *16*. https://doi.org/10.3389/fnbeh.2022.970083

Nyhus, E. (2018). Brain Networks Related to Beta Oscillatory Activity during Episodic Memory Retrieval. *Journal of Cognitive Neuroscience*, *30*(2), 174–187. https://doi.org/10.1162/jocn_a_01194

Nyhus, E., & Curran, T. (2010). Functional role of gamma and theta oscillations in episodic memory. *Neuroscience & Biobehavioral Reviews, Binding Processes: Neurodynamics and Functional Role in Memory and Action*, *34*(7), 1023–1035. https://doi.org/10.1016/j.neubiorev.2009.12.014

Schweinberger, S. R., & Neumann, M. F. (2016). Repetition effects in human ERPs to faces. *Cortex, Special Issue:Repetition Suppression-an Integrative View*, *80*, 141–153. https://doi.org/10.1016/j.cortex.2015.11.001

Sederberg, P. B., Kahana, M. J., Howard, M. W., Donner, E. J., & Madsen, J. R. (2003). Theta and Gamma Oscillations during Encoding Predict Subsequent Recall. *The Journal of Neuroscience*, *23*(34), 10809–10814. https://doi.org/10.1523/JNEUROSCI.23-34-10809.2003

Shirvalkar, P. R., Rapp, P. R., & Shapiro, M. L. (2010). Bidirectional changes to hippocampal theta–gamma comodulation predict memory for recent spatial episodes. *Proceedings of the National Academy of Sciences of the United States of America*, *107*(15), 7054–7059. https://doi.org/10.1073/pnas.0911184107

Summerfield, C., Wyart, V., Mareike Johnen, V., & de Gardelle, V. (2011). Human Scalp Electroencephalography Reveals that Repetition Suppression Varies with Expectation. *Frontiers in Human Neuroscience*, *5*. https://doi.org/10.3389/fnhum.2011.00067

Tamura, M., Spellman, T. J., Rosen, A. M., Gogos, J. A., & Gordon, J. A. (2017). Hippocampal-prefrontal theta-gamma coupling during performance of a spatial working memory task. *Nature Communications*, *8*(1), Article 1.

Tort, A. B. L., Komorowski, R., Eichenbaum, H., & Kopell, N. (2010). Measuring Phase-Amplitude Coupling Between Neuronal Oscillations of Different Frequencies. *Journal of Neurophysiology*, *104*(2), 1195–1210. https://doi.org/10.1152/jn.00106.2010

Vivekananda, U., Bush, D., Bisby, J. A., Baxendale, S., Rodionov, R., Diehl, B., Chowdhury, F. A., McEvoy, A. W., Miserocchi, A., Walker, M. C., & Burgess, N. (2021). Theta power and theta-gamma coupling support long-term spatial memory retrieval. *Hippocampus*, *31*(2), Article 2. https://doi.org/10.1002/hipo.23284

Voytek, B., D'Esposito, M., Crone, N., & Knight, R. T. (2013). A method for event-related phase/amplitude coupling. *NeuroImage*, *64*, 416–424. https://doi.org/10.1016/j.neuroimage.2012.09.023
